# A proximity-dependent biotinylation map of a human cell: an interactive web resource

**DOI:** 10.1101/796391

**Authors:** Christopher D. Go, James D.R. Knight, Archita Rajasekharan, Bhavisha Rathod, Geoffrey G. Hesketh, Kento T. Abe, Ji-Young Youn, Payman Samavarchi-Tehrani, Hui Zhang, Lucie Y. Zhu, Evelyn Popiel, Jean-Philippe Lambert, Étienne Coyaud, Sally W.T. Cheung, Dushyandi Rajendran, Cassandra J. Wong, Hana Antonicka, Laurence Pelletier, Alexander F. Palazzo, Eric A. Shoubridge, Brian Raught, Anne-Claude Gingras

## Abstract

Compartmentalization is a defining characteristic of eukaryotic cells, partitioning cellular processes into discrete subcellular locations. High throughput microscopy^1^ and biochemical fractionation coupled with mass spectrometry^2–6^ have helped to define the proteomes of a variety of organelles and macromolecular structures. However, many other intracellular compartments have remained refractory to such approaches, due for example to difficulty in purifying non-membrane bound structures. Proximity-dependent biotinylation techniques such as BioID provide an alternative approach for defining the composition of cellular compartments in living cells^7–10^. Here we present a BioID-based map of a human cell based on 192 markers from 32 different subcellular compartments, comprising 35,902 high confidence proximity interactions, and defining the intracellular locations of 4,145 unique proteins in HEK 293 cells. Our localization predictions meet or exceed previous-approaches, with higher specificity, and enabled the discovery of proteins at the mitochondrial outer membrane-endoplasmic reticulum (ER) interface that are critical for mitochondrial homeostasis. Based on this dataset, we have established humancellmap.org as a community resource that provides online tools for localization analysis of user BioID data, and demonstrate how this resource can be used to better understand BioID datasets.

---

Proximity-dependent biotinylation approaches are used to characterize the intracellular environment occupied by a protein of interest in living cells^*7,8*^. In the most widely used of these techniques, BioID, a mutant *E. coli* biotin ligase (BirA*, R118G) is fused with the coding sequence of a “bait” polypeptide of interest, and the resulting fusion protein expressed in cultured cells or model organisms^11,12^. The abortive BirA* enzyme generates biotinoyl-AMP, but displays a reduced affinity for this chemical intermediate. Highly reactive biotinoyl-AMP is thus released into the local environment, where it reacts with free epsilon amine groups on lysine residues^7^ within ~10nm of the bait protein^13^. Since biotinylation is covalent, harsh lysis conditions can be used to solubilize proteins from even poorly soluble intracellular compartments (e.g. membranes, chromatin or the nuclear lamina). Biotinylated proteins are then captured with streptavidin (or a streptavidin derivative) linked to a solid phase support, and identified by mass spectrometry (MS). Since the average globular protein of 30–200 kDa is 5–10nm in diameter, the labeling radius of this technique favors the biotinylation of direct binding partners, other components of protein complexes in which the bait resides, and proteins in the immediate intracellular “neighborhood”. BioID has thus been successfully employed by multiple laboratories to define the composition of many different protein complexes, and the spatial organization of a number of both membrane-bound and membraneless organelles (e.g.^7–10^).

To better characterize how the proteome is organized in a human cell, we used BioID to profile 234 previously defined intracellular protein markers for 32 different cellular compartments. These compartments included the cytosolic face of all membrane-bound organelles, the ER lumen, subcompartments of the mitochondria, membraneless organelles such as the centrosome and the nucleolus, and cytoskeletal structures. The endocytic system was also queried to identify components enriched along its continuum (e.g. early versus late endosomes).

Well characterized markers were selected from the literature, with the goal of analyzing multiple independent bait proteins for each subcellular compartment (**Figure 1A; Supplementary Table 1)**. Each compartment marker was tagged with BirA*, stably integrated in HEK293 Flp-In T-REx cells, and processed for BioID (see Methods for details). SAINTexpress^14^ was used to identify high confidence proximity interactors (or “prey” proteins) by scoring spectral counts against a set of negative controls that capture endogenously biotinylated proteins, polypeptides that are non-specifically biotinylated by BirA* and proteins that are non-specifically isolated by the solid phase matrix (sepharose). Only high-confidence interactors (i.e. those passing a 1% FDR threshold) were considered for downstream analysis (**Supplementary Table 2**). Reproducibility was high across replicate analyses of the same marker, with a mean R^2^ of 0.95 (**Supplementary Table 2**). Quality control for each marker also included immunofluorescence microscopy to confirm expected localization (**Figure 1B; Supplementary Table 1)** and Gene Ontology (GO) enrichment analysis of the identified high confidence prey proteins, to ensure enrichment of expected cellular component (CC) terms (**Supplementary Table 3**). With the notable exception of the Golgi lumen (for which all tested baits remained trapped in the ER), all selected compartments were successfully characterized with multiple baits.

**Fig. 1:**
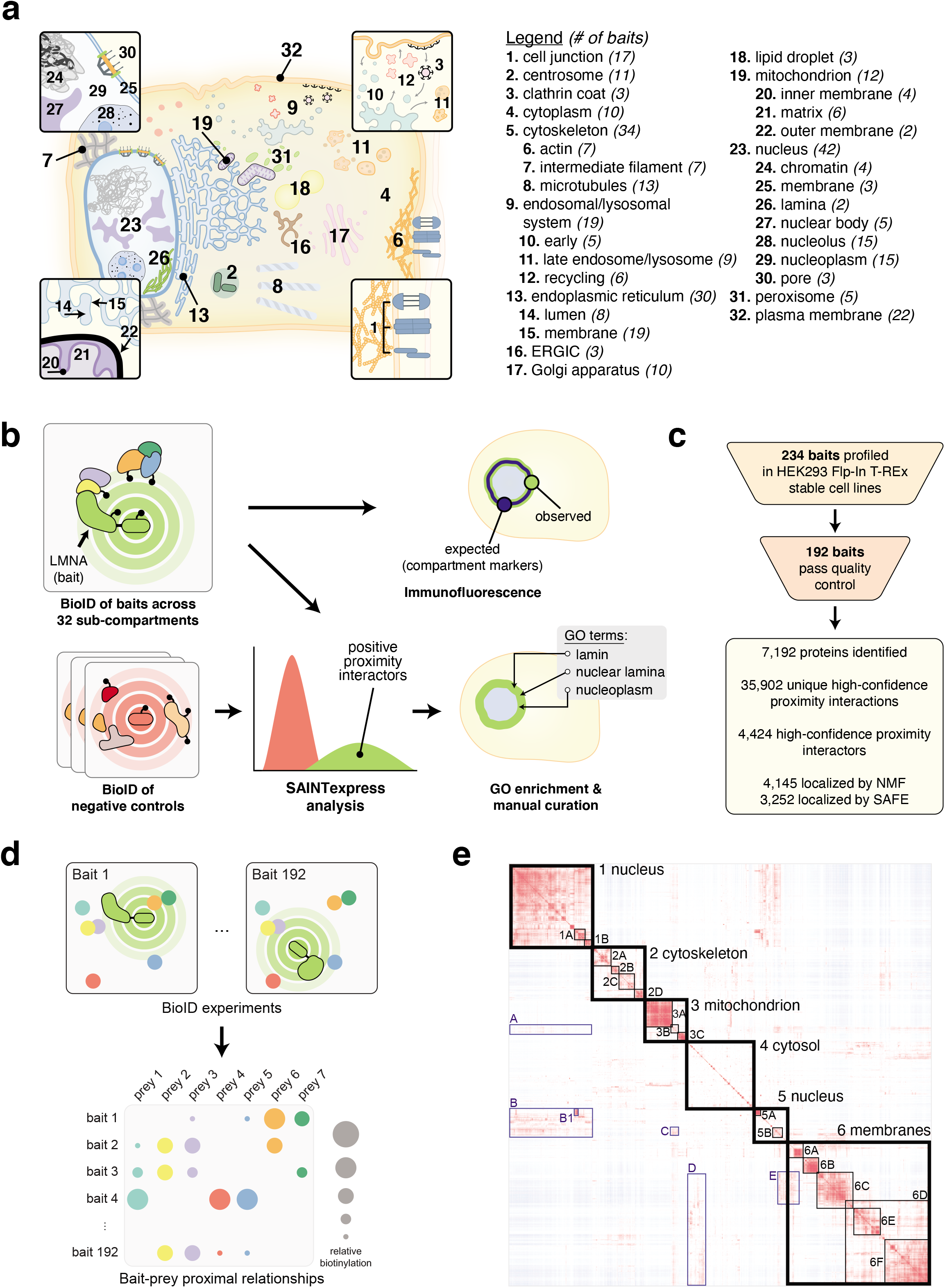
Dataset generation and localization rationale. (**a**) Cellular compartments targeted for profiling by BioID. **Bold** numbers on the schematic correspond to the indices in the legend. *Italicized* numbers in brackets next to the compartment name indicate the number of baits used to profile that compartment after quality control (QC). (**b**) QC and dataset-scoring pipeline. Bait localization was assessed by immunofluorescence (IF) alongside endogenous compartment markers, and GO term enrichment of significant proximity interactors following SAINTexpress^14^ scoring against control replicates. All IF images are available with their corresponding bait reports at humancellmap.org. **(c)** QC excluded 42 baits from the original set of 234, and SAINT analysis of the final bait set yielded 35,902 interactions with 4,424 unique high confidence proximity interactors. (**d**) Rationale for prey-association based localization. In BioID, bait proteins label the proximal environment in a distance-dependent manner. The relative labelling of preys across baits is therefore dependent on the proximity of those prey proteins to each other *in situ.* In other words, each prey produces a “signature” across baits that it will share with proteins localizing to the same locales. This correlation can be used to assign localizations to preys based on the previously known localization of preys with a similar signature. (**e**) Correlation prey-prey heat map. Correlation between preys across baits was calculated from spectral counts using the Pearson coefficient, and preys were clustered by Euclidean distance and complete linkage. The heat map was manually annotated by performing GO enrichment on cluster components. See **Supplementary Table 6** for annotations and GO enrichments of the highlighted clusters. Interestingly, there are many subclusters within large organelle clusters that are much bigger than traditional protein complexes, suggestive of a layer of organization between complexes and the organelle, possibly akin to protein communities^44^.

192 candidate markers passed quality control (**Figure 1C**), establishing 35,902 interactions with 4,424 unique high confidence proximity interactors. BioID bait profiles were overall consistent with their expected compartments (**Extended Data Figure 1**), further attesting to the quality of the dataset. As expected from previous studies^8,15^, bait proteins that profiled different faces of the same organelle yielded distinct proximity profiles, since biotin labelling is restricted to the membrane surface exposed to BirA* enzymatic activity. For example, using the Jaccard index as a measure of similarity (where 1 indicates complete overlap, and 0 indicates no overlap), we observed a median Jaccard index of 0.541 amongst the eight baits used to profile the ER lumen. A much smaller median Jaccard index of 0.069 was observed between the interactomes of baits localized to the ER lumen versus 19 ER membrane-bound cytosol-facing baits. Similarly, in the mitochondria the median Jaccard index for the five matrix baits is 0.587, while the median value between the matrix and six non-matrix mitochondrial baits is 0.065.

Consistent with BioID being a proximity-dependent labelling technique, a clear correlation was observed between prey abundance (length normalized spectral counts) observed by MS and the probability that a prey protein was previously reported as an interactor for a given bait (based on BioGRID^16^ and IntAct^17^ protein interaction databases; **Extended Data Figure 2A-B**; **Supplementary Table 4**). Protein turnover rate^18^ and the number of lysines per protein did not appear to be correlated with prey detection, but protein expression level was^19^ (**Extended Data Figure 2C-F**; **Supplementary Table 5**). While baits localizing to the same compartment had similar prey profiles, they did not display perfect overlap. For example, in the mitochondrial matrix, the tRNA synthetase AARS2 and pyruvate dehydrogenase PDHA1 interactomes share a Jaccard Index of 0.66, but the AARS2 interactome is preferentially enriched for components of the mitochondrial ribosome and proteins involved in translation, while PDHA1 preferentially recovers pyruvate dehydrogenase complex components (**Extended Data Figure 3**). These results are expected, based on the known interactions and functions of AARS2 and PDHA1, and highly reproducible (**Supplementary Table 2**). As also discussed elsewhere^9,10,20,21^ BioID can thus provide both compartment and sub-compartmental resolution, uncovering distinct protein environments within the same cellular compartment.

To localize preys to discrete intracellular subcompartments, we exploited the fact that prey proteins with correlated behavior, i.e. those that interact with the same set of baits, are likely to reside in the same multiprotein complex, organelle or subcellular region^10^ (**Figure 1D**). Analysing the data using a prey-centric approach also ensures robustness across the dataset to unexpected bait behavior (e.g. subtle mislocalization associated with tagging and expression, localization to more than one compartment, etc.). Here, we first calculated the Pearson correlation of prey profiles, which highlighted a number of clearly defined (sub)compartments (**Figure 1E**; see **Supplementary Table 6** for cluster annotation). This was followed by Spatial Analysis of Functional Enrichment (SAFE)^22^ to annotate correlated preys (see **Figure 2A** and Methods for details), which localized 3,252 of the 4,424 high confidence prey proteins to 23 different intracellular compartments (**Extended Data Figure 4; Supplementary Table 7)**. In addition to annotation based on correlation, we also applied Non-Negative Matrix factorization (NMF)^23^, which localized 4,145 preys to 20 compartments (**Figure 2A,B, Extended Data Figure 5; Supplementary Table 8)**. An advantage of NMF is that it can score prey proteins across multiple subcellular locations, enabling the assignment of localizations to more than one locale (discussed more below). 54% of the localizations assigned by SAFE were previously reported, and 50% for NMF (based on the highest scoring NMF compartment for each prey). When both SAFE and NMF made a prediction (3,252 proteins), they were consistent in 88% (2,855) of cases (**Supplementary Table 9; Supplementary Table 10**). While many subcompartments, such as the ER membrane (**Figure 2B)** versus ER lumen; the mitochondrial matrix, inner and outer membranes; and chromatin, nucleolus, nuclear body and nucleoplasm, were effectively separated, a lack of resolving power was evident for subcompartments associated with (endo)membranes. This result is consistent with the fluid nature of these compartments, and bait localization that spans multiple vesicle subpopulations. Accurately defining profiles for these more challenging subcompartments will require additional study.

**Fig. 2:**
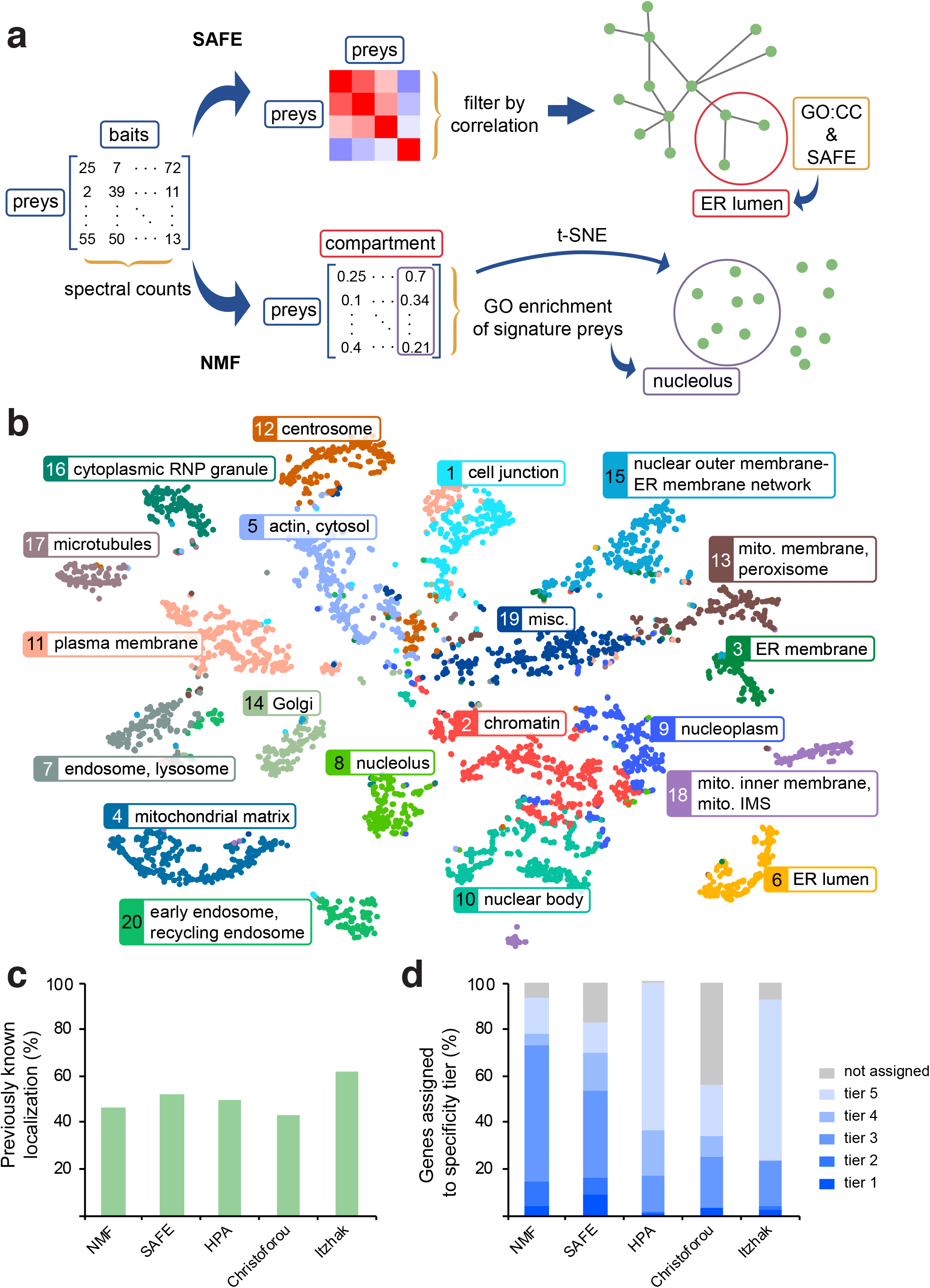
Localization of proteins using prey-prey correlation information. (**a**) Pipelines for localizing prey proteins using Spatial Analysis of Functional Enrichment (SAFE^22^) and Nonnegative Matrix Factorization (NMF^23^). In SAFE, preys with a correlation across baits ≥ 0.65 are considered “interactors” and these pairs are used to generate a network that is annotated for GO:CC terms (refer to methods). In NMF, the bait-prey spectral count matrix is reduced to a compartment-prey matrix and compartments are then defined using GO:CC for the compartment’s most abundant preys. A 2D network is generated in parallel from the compartment-prey matrix using t-SNE^45^. (**b**) NMF-based map of the cell generated with t-SNE. Each prey is coloured to indicate its primary localization. An interactive version of the map can be viewed at humancellmap.org/explore/maps. (**c**) Performance of the NMF- and SAFE-based procedures against the immunofluorescence-based Human Protein Atlas (HPA^1^, www.proteinatlas.org) and the fractionation studies of Christoforou^2^ and Itzhak^3^. The vertical axis indicates the number of genes assigned to a previously known localization (GO:CC term). (**d**) Specificity of localization assignments. GO:CC terms were binned into specificity tiers (1: most specific; 5: least specific), and the number of genes assigned to each tier was quantified for our pipelines and the published studies under comparison.

Compartments identified by the NMF and SAFE pipelines displayed enrichment for expected protein domains and motifs (**Supplementary Table 11**). For example, the plasma membrane compartment is significantly enriched for pleckstrin homology (PH), immunoglobulin, RhoGAP, RhoGEF and tyrosine kinase domains, while the related cell junction compartment is enriched for PDZ and FERM domains. Nuclear subcompartments are similarly distinguished, with the chromatin compartment enriched for the KRAB domain, C2H2 zinc fingers, bromodomains and PWWP domain; the nucleolus for the DEAD and helicase domains; and the nuclear body compartment for RNA recognition motif (RRM) and G-patch domains. Almost no enriched domains were shared between compartments. Compartments also displayed clear enrichment for specific protein sequence motifs or sequence characteristics (coiled-coiled, disordered, low complexity, signal peptide and transmembrane), but in contrast with domains, sequence motifs were often shared between compartments (**Extended Data Figure 4B; Extended Data Figure 5B**).

We next compared our localization predictions with those made by several large-scale microscopy and fractionation studies that were similar in scope and generally aimed at targeting the same organelles. After removing Human Protein Atlas (HPA^1^) annotations from GO to prevent self-validation, our recovery of known protein localizations for NMF and SAFE analyses was similar to HPA and fractionation approaches (**Figure 2C; Supplementary Table 12**). However, this simple analysis ignores the fact that less specific localization annotations (e.g. nucleus versus cytoplasm) are more likely to be known for a protein than more specific subcompartment localizations (**Extended Data Figure 6A**). By binning GO localization annotations into “precision tiers” based on information content (see Methods for details), with tier 1 containing the most specific localizations (e.g. terms such as “peroxisome” or “spliceosome”) and tier 5 the least specific (e.g. “cytoplasm” or “nucleus”), we found that our predictions are actually more specific than those of other approaches. For example, 73% of proteins were localized to the tier 3 bin or better in our NMF analysis (54% for SAFE), versus 17-25% for the other data sets (**Figure 2D; Supplementary Table 12**). High recall of known localizations with increased specificity is thus a marked advantage of our methodology (**Extended Data Figure 6B and C**). As the human cell map project expands in the future through the incorporation of additional baits, we expect to be able to generate even more precise localization predictions.

To further assess the accuracy of our predictions, we used immunofluorescence (IF) microscopy, focusing on proteins without a clear annotation (e.g. annotated as an Open Reading Frame, ORF, or simply as “family with sequence similarity”, FAM) and protein families that share domains or structures, but with different predicted localizations. These included proteins annotated as solute carriers (SLC), transmembrane proteins (TMEM), and the Rab family of small GTPases. We also only selected proteins for which the NMF and SAFE predictions were in agreement. 65 GFP-tagged prey proteins that were assigned the same annotations by NMF and SAFE were transiently expressed in HEK293 cells, and colocalization with well characterized markers was assessed. 86% (56/65) of the predictions tested were supported by this method (**Figure 3A, Extended Data Figure 7; Supplementary Table 13**). Twenty of the proteins validated by transient expression were also stably expressed in HEK293 cells, with 17 recapitulating the initial validation (**Supplementary Table 13**). Together, these results demonstrate the ability of our analysis pipeline to correctly assign subcellular localization to poorly characterized proteins.

**Fig. 3:**
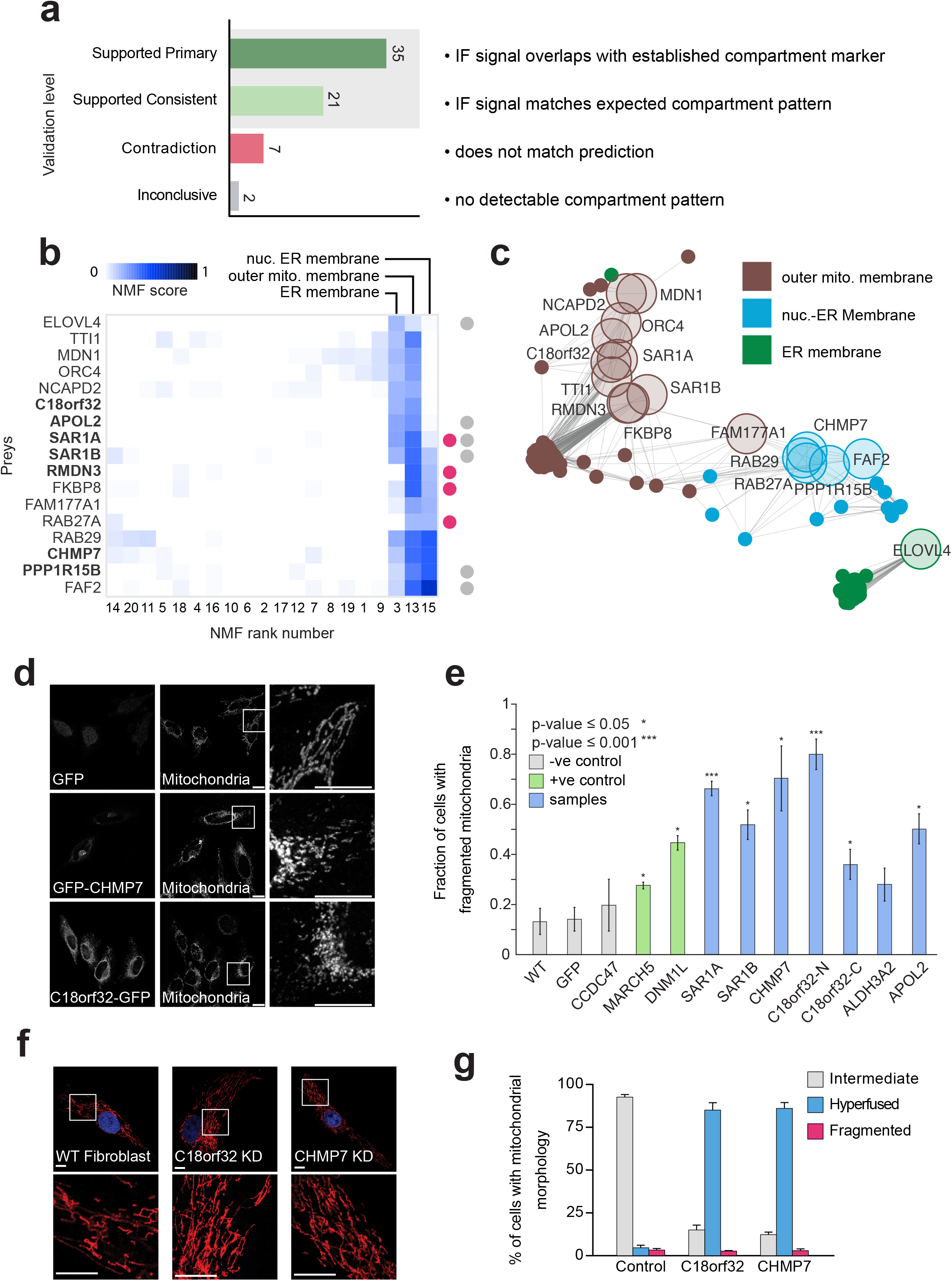
Novel protein localizations and identification of proteins at mitochondrial-ER contact sites. (**a**) Summary of experimental validations for predicted localizations of proteins by immunofluorescence (IF) microscopy. Confidence rankings were annotated as follows: “supported primary” indicates proteins that matched the NMF and SAFE prediction; “supported consistent” indicates proteins that matched the NMF and SAFE prediction, but did not have an endogenous compartment marker for the immunofluorescence microscopy; “contradiction” indicates proteins that failed to localize to the predicted localization made by NMF and SAFE; “inconclusive” indicates proteins that had no clear subcellular compartment localization. (**b**) Heat map of genes with a primary localization at the mitochondrial outer membrane or the ER membrane/nuclear outer membrane-ER membrane and a secondary localization to the other compartment as computed by NMF. To be included on the heat map, genes required an NMF score of at least 0.15 in the compartments of interest, a score ratio of at least 0.4 between the primary and secondary localization, and a score ratio of at least 2 between the compartments of interest and all other compartments. **Bold** genes indicate those selected for mitochondrial morphology assays in the following panels. A grey-dot on the right side of the plot indicates proteins involved in lipid and cholesterol homeostasis, while a pink dot indicates calcium signalling. (**c**) Sub-network from **Extended Data Figure 5A** of the ER-mito proteins selected for follow-up and their first degree neighbors. ER-mito proteins are shown as large transparent nodes with labels. First-degree neighbors are shown as small filled nodes. Edges represent proteins with a correlation score ≥ 0.9 across all NMF compartments. Green nodes indicated proteins localized to the ER membrane, blue nodes indicate proteins localized to the nuclear-ER membrane and brown nodes indicate proteins localized to the mito-peroxisome membrane. (**d**) Mitochondrial morphology is altered by transient expression of GFP-tagged CHMP7 and C18orf32, as monitored by confocal IF microscopy in HeLa cells. Cells were fixed and probed with antibodies directed against GFP and COX IV (see Methods for details). The white box indicates the zoomed area displayed in the rightmost panels. Scale bars, 10 μm. **(e)** Quantification of HeLa cells with fragmented mitochondrial morphology upon overexpression of GFP-tagged proteins. Negative controls are coloured in grey, including untransfected cells, cells expressing GFP alone and cells expressing CCDC47 (an ER protein); positive controls MARCH5 and DNM1L are coloured in green; test candidates are coloured in blue. Experiments were done in biological triplicate with an average of ~150 cells counted per sample, statistical confidence of mitochondrial fragmentation was calculated using the Student’s t-test and error bars represent standard deviation (see Methods). **(f)** IF of mitochondrial morphology in human primary fibroblasts in which GFP, CHMP7 or C18orf32 were targeted by siRNA-mediated knockdown. The white box indicates the zoomed area displayed in the bottom panels. The mitochondrial marker is an antibody against cytochrome C (see Methods). Scale bars, 10 μm. **(g)** Quantification of primary fibroblast mitochondrial morphology upon siRNA-mediated knockdown. Fraction of cells with hyperfused, fragmented and intermediate mitochondrial morphology are displayed in blue, red and grey respectively. Experiments were done in biological triplicate with 100-150 cells counted per sample and error bars represent standard deviation (see Methods).

While BirA* localized to two different faces of the same intracellular membrane generates distinct interactomes, we also found that the topology of membrane-associated prey proteins could be predicted from our data, based on the exposure of non-transmembrane stretches of protein sequence to the cytosol or lumen (**Extended Data Figure 8A**). For example, all transmembrane-domain-containing prey proteins localized to the ER membrane were assigned a cytosol/lumen ratio (CLR) score, based on their NMF localization scores for the cytosolic or lumenal faces of the ER. For proteins with known sequence orientation information, this metric displayed clear correlation (R^2^ = 0.42) with existing annotations (**Extended Data Figure 8B,C and Supplementary Table 14)**. Not all preys in this analysis have a previously assigned orientation (i.e. unknown orientation of protein N- and/or C-termini toward the lumen or cytosol), allowing these scores to be used as predictions for follow-up studies.

While many proteins (684) yielded data consistent with exchange between contiguous intracellular compartments (e.g. between the cell junction and plasma membrane; early and late endosomes; and across nuclear substructures), only 305 (7%) prey proteins displayed a clear signature in at least two non-contiguous cell compartments (**Supplementary Tables 8 and 15**), a phenomenon that could be due to moonlighting^1,24^ or localization to membrane contact sites^25^. We also examined the degree of intra- and inter-compartmental crosstalk between compartments but found very little taking place between non-contiguous compartments (**Extended Data Figure 9, Supplementary Table 15**). The lack of widespread moonlighting behavior or inter-compartmental interactions in our dataset could be due to the relatively long biotinylation times used in the classic BioID approach used here, which could disfavor more dynamic and/or condition-dependent localizations. Future experiments using faster enzymes^26^ (which provide results comparible with the original BirA* enzyme used here; **Extended Data Figure 10, Supplementary Table 16**) may help to better define moonlighting activities.

Contacts between the mitochondria and ER are critical for lipid and calcium exchange, and play important roles in mitochondrial dynamics^27,28,29^. Moonlighting and inter-compartmental edge analyses identified 17 proteins linked to both the mitochondria/peroxisome and ER membrane compartments (**Figure 3B-C, Supplementary Table 17**), including SAR1A and SAR1B, two GTPases that regulate mitochondria-ER contact sites^30^, and RMDN3 (aka PTPIP51), a mitochondria-ER tether protein that regulates autophagy through calcium signaling^31,32^. We prioritized proteins from this list with an ER annotation in the literature for follow-up (SAR1A, SAR1B, APOL2, C18orf32, CHMP7 and PPP1R15B), and performed BioID to localize them more precisely. While strongly enriching for expected ER components, SAR1A, SAR1B, C18orf32 and CHMP7 (and to a lesser extent APOL2 and PPP1R15B) also recovered outer mitochondrial membrane proteins such as AKAP1, MAVS, HK1 and OCIAD1, as well as other predicted ER-mitochondrial contact site candidates (**Extended Data Figure 11; Supplementary Table 17**). This set of baits also detected two proteins linked to mitochondrial fission, DNM1L (orthologous to yeast Drp1^33^) and INF2 (a formin that mediates actin-dependent fission^34^). To test whether the proteins selected for follow-up could play a role in mitochondrial dynamics, we first expressed GFP-tagged versions of these proteins and quantified the percentage of cells with fragmented mitochondrial morphology. In this assay, MARCH5/MITOL (a ubiquitin E3 ligase that positively regulates membrane fission^35^) and DNM1L served as positive controls, inducing mitochondrial fragmentation as expected (**Figure 3D, 3E**). Expression of three of our candidate proteins, APOL2 (apolipoprotein L2), C18orf32 (a protein that traffics to lipid droplets^36^) and CHMP7 (an ESCRT-III component) also strongly induced mitochondrial fragmentation. Conversely, siRNA-mediated knockdown of C18orf32 and CHMP7 induced a striking hyperfused mitochondrial phenotype, suggesting that these proteins are important for mitochondrial homeostasis (**Figure 3F, 3G**). While their specific roles in mitochondrial dynamics remain to be defined, these data demonstrate how our data can be mined to reveal novel protein functions within and between different subcellular compartments.

To facilitate exploration of our dataset, and to create a repository for BioID data generated by our lab and collaborators moving forward, we created humancellmap.org. Here the community can search and view data on all profiled baits, identified preys, compartments and organelles in this project, and explore interactive 2D maps and networks of NMF and SAFE data. “Help” documentation describing all available features for the site can be found at humancellmap.org/help. The site will be updated as new data become available.

An additional key feature of the website is the ability to upload user BioID data and compare it to the humancellmap database (**Figure 4**; humancellmap.org/query). This analysis can help to localize a query bait to a specific subcellular compartment(s) based on interactome similarity with other baits in the dataset, identify previously queried baits with similar interactomes, and identify those preys that are most specific to the queried bait. To illustrate these functions, we performed BioID on a regulatory subunit of PI3 kinase (PIK3R1), an SH2 domain-containing adaptor that recruits PI3 kinase to activated receptor complexes at the plasma membrane. Analyzing its BioID data using humancellmap.org revealed that the baits in our dataset most similar to PIK3R1 localize to the plasma membrane and cell junction. Importantly, while 27 high-confidence proximity interactions were detected with this bait, a specificity metric revealed highly specific proximity interactions with PI3 kinase catalytic subunits, insulin receptor substrate proteins (IRS2, IRS4) and the scaffold protein GRB2^37^ (**Figure 4A, B**). We also reanalyzed a previously published BioID bait, RNGTT^10^, a nuclear protein involved in mRNA capping. Humancellmap analysis reported a nuclear localization, with bait-specific interactions including several RNA Polymerase II subunits and components of the catalytic subunit of the PP4 phosphatase, as previously reported^38,39^ (**Figure 4C**). This strategy can also be expanded to the analysis of poorly characterized proteins (**Supplementary Table 18**). FAM171A1 and FAM171B were predicted by our NMF and SAFE analyses to localize to the cell junction and plasma membrane. Consistent with this prediction, their BioID profiles when screened as baits were most similar to junctional and plasma membrane baits, while bait-specific preys included several cytoskeletal proteins, in line with a previous study that reported a reduction of actin stress fibers following knockdown of FAM171A1^40^ (**Extended Data Figure 12A**). MTFR2 (FAM54A) was associated with the mitochondrial outer membrane and peroxisome as a prey protein, with a weak signature at the mitochondrial inner membrane/mitochondrial intermembrane space. When profiled and screened as a bait, the analysis module reports that it is most similar to peroxisomal baits, followed by mitochondrial outer and inner membrane baits, supporting its predicted localization. Interactions with MTFR1, SLC25A46 and VPS13D were found to be highly specific to MTFR2, consistent with the mitochondrial fragmentation previously observed upon overexpression of GFP-MTFR2^41^ (**Extended Data Figure 12B**). Finally, we re-analyzed BioID data on the bromodomain-containing protein BRD3 in cells treated with the BET inhibitor JQ1, which leads to a relocalization of BRD3 to the nucleolus^42^. This relocalization was apparent when analyzed against the humancellmap, with JQ1-treated BRD3 more similar to nucleolar baits (**Extended Data Figure 13**, **Supplementary Table 19**), attesting to the applicability of the humancellmap.org resource for the exploration of conditiondependent BioID datasets.

**Fig. 4:**
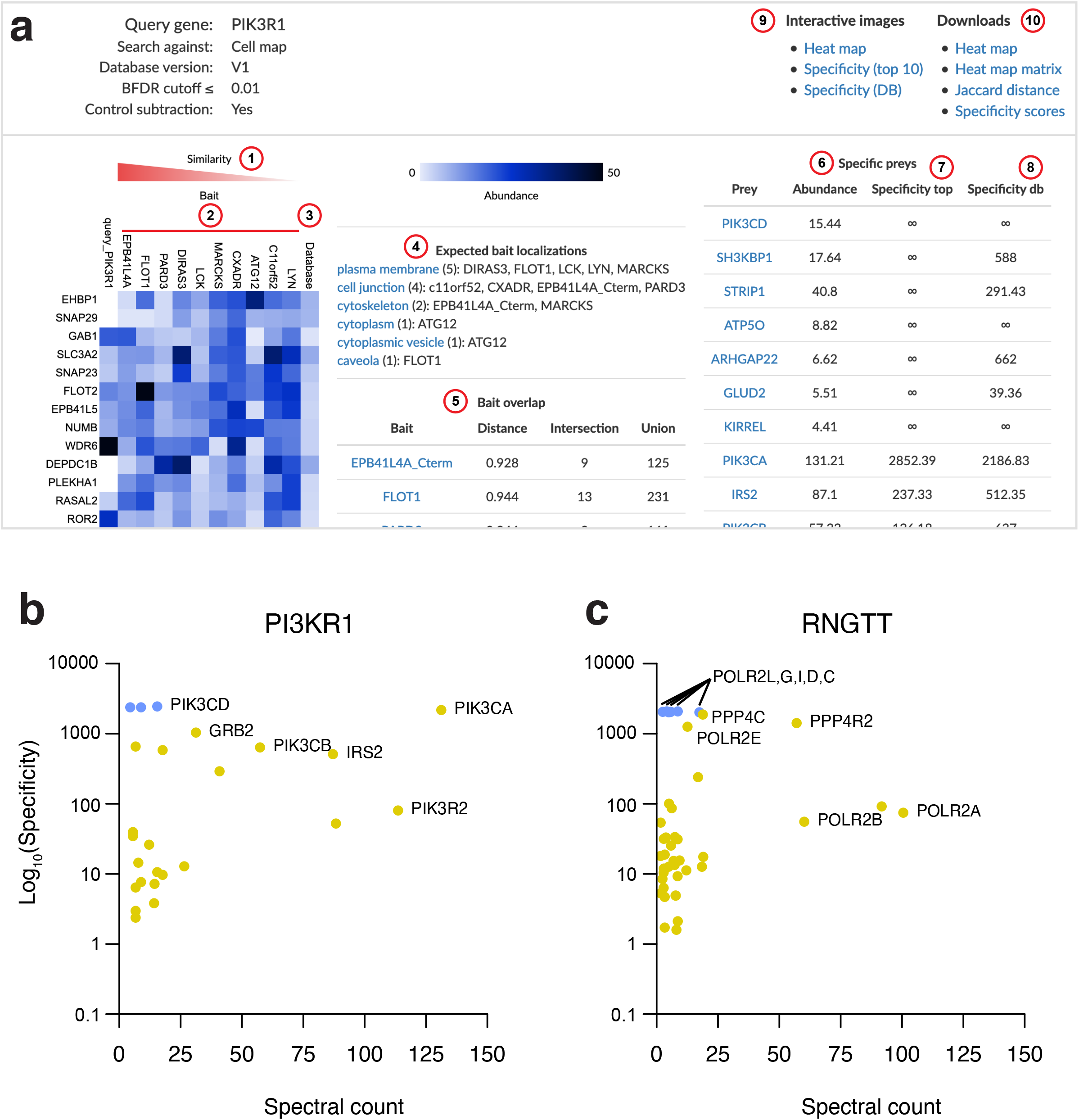
Analysis module at humancellmap.org. **a**) Screenshot of the analysis report for the bait PIK3R1. Red circles indicate: 1) Baits from the humancellmap are sorted from most similar to least similar as calculated by the Jaccard distance. 2) The ten most similar baits to the query in the humancellmap. 3) The average spectral count for each prey averaged across all baits in the humancellmap database. 4) Expected localizations of the ten most similar baits. 5) Overlap/similarity metrics between the query bait and the top ten most similar baits in the humancellmap. The distance is the Jaccard distance, with a score of 0 for complete prey overlap and 1 for no overlap. The intersection refers to the number of shared preys, and the union refers to the combined number of preys between the query and the indicated bait. 6) The most specific preys for the query. The specificity score is calculated as the fold enrichment of a prey in the query relative to the average across the humancellmap baits used for the comparison. 7) The specificity score calculated against the top ten most similar baits to the query. 8) The specificity score calculated against all baits in the humancellmap. 9) Links to open the heat map or specificity plots at the interactive viewer at ProHits-viz^46^. 10) Links for data downloads. **b**) Specificity plot for PIK3R1 showing the control subtracted spectral counts versus the specificity score (see **Methods** for specificity score calculation). **c**) Specificity plot for RNGTT.

While the current version of humancellmap.org is a static view of a single cell type (HEK293), future versions will explore higher density coverage of baits (by e.g. merging organelle-specific datasets^9,10,20^ into humancellmap.org), include other cell types, and feature dynamic interactomes profiled with faster enzymes, thereby supplementing other proteomics and cell biological resources.

## Supporting information

Website help documentation

## Acknowledgements

We thank Zhen-Yuan Lin for profiling PIK3R1 and members of the Gingras lab for helpful discussion and advice throughout the project, and Jin Zhang and many cell biologists for helpful suggestions regarding bait selection.

Work in the Gingras lab was supported by a Canadian Institutes of Health Research (CIHR) Foundation Grant (FDN 143301). EAS is supported by a grant from the CIHR (MOP-133530). Proteomics work was performed at the Network Biology Collaborative Centre at the Lunenfeld-Tanenbaum Research Institute, a facility supported by Canada Foundation for Innovation funding, by the Ontario Government, and by Genome Canada and Ontario Genomics (OGI-139). This research was enabled in part by support provided by Compute Canada (www.computecanada.ca). C.D.G. was supported by a CIHR Banting studentship. A.-C.G. is the Canada Research Chair in Functional Proteomics and the Lea Reichmann Chair in Cancer Proteomics.

## Data availability

Datasets consisting of raw files and associated peak lists and results files have been deposited in ProteomeXchange through partner MassIVE (http://proteomics.ucsd.edu/ProteoSAFe/datasets.jsp) as complete submissions. Additional files include the sample description, the peptide/protein evidence and the complete SAINTexpress output for each dataset, as well as a “README” file that describes the dataset composition and the experimental procedures associated with each submission. The different datasets generated here were submitted as independent entries.

Dataset 1 (see **Supplementary Table 2**):

Go_BioID_humancellmap_HEK293_lowSDS_core_dataset_2019

MassIVE **ID** MSV000084359 and PXD015530

Dataset 2 (see **Supplementary Table 2**): Go_BioID_humancellmap_HEK293_ highSDS_core_dataset _2019

MassIVE ID MSV000084360 and PXD015531

Dataset 3 (see **Supplementary Table 18**): Go_BioID_humancellmap_HEK293_prediction_2019 MassIVE ID MSV000084369 and PXD015554

Dataset 4 (see **Supplementary Table 17**): Go_BioID_humancellmap_HEK293_ER-mito_candidates_2019

MassIVE ID MSV000084357 and PXD015528

Negative control samples were deposited in the Contaminant Repository for Affinity Purification^43^ (CRAPome.org) and assigned samples numbers CC1100 to CC1185 (see **Supplementary Table 2**); this will be part of the next release of the database.

## Code Availability

Source code used for analysis can be accessed from github.com/knightjdr/cellmap-scripts.

## Material Availability

All DNA vectors used in this study can be obtained by contacting the corresponding author.

## Author Contributions

A.-C.G., C.D.G. and J.D.R.K. conceived the project.

A.-C.G., C.D.G., J.D.R.K. and B.R. wrote the paper with input from G.G.H, P.S.T, J.-Y.Y., J.-P.L and E.C.

C.D.G. generated most of the BioID constructs and cell lines and performed BioID experiments and immunofluorescence studies.

J.D.R.K., C.D.G., G.G.H. and A.-C. G. performed data analysis.

J. D.R.K. created the humancellmap.org website.

K. A. contributed the cell model and illustrations.

A. R., H.A. and E.S performed mitochondrial morphology experiments and analyzed results.

B. R. generated constructs and cell lines for BioID and testing predictions.

G.G.H., K.A., J.-Y.Y., P.S.-T., H.Z., L.Y.Z., E.P., J.-P.L., E.C., S.C., L.P., B.R. and A.P. contributed constructs and cell lines.

A.-C.G. supervised the project.

---

**Extended Data Figure 1**: Bait similarity and localization. The Jaccard index was calculated between each pair of baits in the core dataset using the list of high confidence (1% FDR) interactors. Baits were clustered using the Euclidean distance and complete linkage method, and clusters optimized using the CBA package in R. The color gradient next to the bait labels indicates whether a bait shares an expected localization with both adjacent baits (red), one adjacent bait (light red) or neither adjacent bait (white). Major clusters were manually annotated based on expected localization of the components.

**Extended Data Figure 2**: Recovery of known preys and factors affecting prey labelling. For each bait the relative proximity of every prey (proximity order) was calculated from the control-subtracted length-adjusted spectral count (CLSC) as described in the **Methods**, such that the prey with the highest CLSC value was considered to be the “interactor” most proximal to the bait and the lowest CLSC value the most distal. **a**) After sorting preys by proximity order and grouping by order across baits, the proportion of previously reported preys was calculated for the *n*^th^ proximity order for *n* = 1, 2,.. 200. **b**) Number of baits with a minimum of *n* preys at a 1% FDR, for *n* = 1, 2,. 200. **c**) Proximity order vs protein turnover rate (hours) in HeLa cells (turnover data is from ^18^). **d**) Proximity order vs protein expression as represented by the log10 normalized MS1 iBAQ intensity from ProteomicsDB^19^. **e**) Proximity order vs the number of lysines per protein. **f**) Log10 normalized MS1 iBAQ intensity of proteins expressed in HEK293 versus HeLa cells from ProteomicsDB^19^. The similarity in proteomes supports the usage of HeLa data in panel **c** as suitable HEK293 data was not available. Values along the x-axis could reflect zero expression or missing data in HeLa cells. These were ignored when calculating the R^2^.

**Extended Data Figure 3**: Bait comparisons for a pair of mitochondrial matrix proteins. Control-subtracted spectral counts are plotted for all high confidence preys (1% FDR) detected with either bait pair under comparison. AARS2 preferentially enriches components of the mitochondrial ribosome and proteins involved in translation, such as GFM1, MRPS9 and TRMT10C, while PDHA1 preferentially interacts with the pyruvate dehydrogenase complex component DLAT and the mitochondrial membrane ATP synthase ATP5F1B.

**Extended Data Figure 4**: SAFE-based map of the cell and motif enrichment. **a**) SAFE-based map of the cell generated from preys with a Pearson correlation score of 0.65 or higher and plotted using Cytoscape with a spring-embedded layout. Each prey is coloured to indicate its primary localization (domain in SAFE terminology) as indicated in the legend. An interactive version of the map can be viewed at humancellmap.org/explore/maps and toggling from NMF to SAFE on the bottom menu. **b**) Pfam regions/motifs enriched in the indicated SAFE domains. The heat map value represents the log2 fold-change between the genes localized to the rank and all preys in the dataset. Only compartments/domains with a significant fold change for at least one motif are displayed on the heat map.

**Extended Data Figure 5:** NMF-based correlation map of the cell and motif enrichment. **a**) NMF-based map of the cell generated from preys with a Pearson correlation score across NMF ranks of 0.9 or higher and plotted using Cytoscape with a spring-embedded layout. Each prey is coloured to indicate its primary localization (rank in NMF terminology) as indicated in the legend. An interactive version of the map can be viewed at humancellmap.org/explore/maps and toggling from t-SNE to correlation on the bottom menu. **b**) Pfam regions/motifs enriched in the indicated NMF ranks. The heat map value represents the log2 fold-change between the genes localized to the rank and all preys in the dataset. Only compartments/ranks with a significant fold change for at least one motif are displayed on the heat map.

**Extended Data Figure 6**: Localization benchmarking. **a**) Percentage of genes localized to a previously known compartment for each specificity tier using our NMF and SAFE pipelines, compared with the Human Protein Atlas (HPA^1^, www.proteinatlas.org) and the fractionation studies of Christoforou^2^ and Itzhak^3^. Specificity tiers were defined by binning GO:CC terms based on their Information Content (IC) as defined in **Methods**. Tier 1 terms are the most specific and Tier 5 the least specific. **b-c**) Percentage of preys localized to a previously known compartment relative to the number of baits they were detected with for NMF and SAFE respectively. **c-d**) Percentage of preys localized to a previously known compartment relative to the average number of spectral counts they were seen with for NMF and SAFE. Preys were binned by spectral count. The left tick mark for each data point indicates the lower bound for the bin (inclusive) and the right tick mark the upper bound (exclusive).

**Extended Data Figure 7**: Localization prediction validation strategy and examples. Confidence rankings are defined as follows: “supported primary” indicates proteins that matched the NMF and SAFE prediction; “supported consistent” indicates proteins that matched the NMF and SAFE prediction but did not have an endogenous compartment marker for the immunofluorescence microscopy; “contradiction” indicates proteins that failed to localize to the predicted localization made by NMF and SAFE; “inconclusive” indicates proteins that had no clear subcellular compartment localization. Representative IF images are shown, markers used are on the respective panel. NMF scores across the defined ranks/categories/compartments are displayed as seen on humancellmap.org with the highest NMF category corresponding to the localization prediction.

**Extended Data Figure 8**: Predicted vs annotated proportion of protein exposed to the cytosol or lumen for ER transmembrane proteins. **a**) Hypothetical examples of proteins with varying proportions of their sequence exposed to the cytosol or lumen. The extent of labelling by cytosolic or lumenal baits should be directly related to proportion of the sequence, and hence lysines available for biotinylation, exposed to the respective faces of the membrane that the protein spans. **b**) All transmembrane domain containing prey proteins localized to the cytosolic face of the ER (NMF compartments 3 and 15) and the lumenal face (NMF compartment 6), were assigned a *cytosol-lumen ratio (CLR) score* based on their NMF profile and sequence data (313 proteins). A prey’s CLR score is calculated by taking the score in the cytosolic facing compartment / maximum score in that compartment and subtracting the corresponding score in the lumenal compartment. A score closer to 1 would indicate a protein with a signature at the cytosolic face of the ER membrane but little or no signature in the lumen and a score of −1 would indicate the opposite. A similar sequence-based score was calculated as the fraction of the sequence annotated as cytosolic minus the fraction that is lumenal according to UniProt. KTN1 is mis-annotated in UniProt^47^ and should have a sequence score of +0.9742. **c**) Three example of proteins and their topology, CLR and sequence scores. Green examples have predictions matching annotated topology.

**Extended Data Figure 9**: Moonlighting and connections between compartments. **a**) Primary and secondary localizations of moonlighting preys. Preys with a score of at least 0.15 in each of two non-contiguous NMF compartments were considered to moonlight (see **Supplementary Table 15** for the list of non-contiguous compartments). The number of preys with a primary localization defined on the vertical axis and a secondary localization defined on the horizontal axis is shown (maximum 18). **b**) Inter-compartment edges were counted for each NMF rank/compartment. An interaction edge was defined between prey pairs having a correlation score across all NMF compartments of at least 0.9. Edges where then defined as “intra-compartment” (if the primary localization for the two preys was the same compartment) or “inter-compartment” (if the primary localization for the two preys was in different compartments) (**Supplementary Table 15)**. Most organelles displayed a much greater proportion of intra-compartment interactions, with the extreme case of the mitochondrial matrix having only 15 inter-compartmental edges out of a total of 37,387 edges. The proportion of inter-compartment edges from the source to each target compartment is shown here. Intercompartmental edges generally conformed with expectations, for example with edges from the chromatin compartment connecting to other nuclear substructures with which they may exchange components. The NMF rank number is shown in brackets next to the source compartment name.

**Extended Data Figure 10**: Comparison of prey profiles for LMNA tagged with BioID, miniTurbo and TurboID. **a**) Spectral counts for significant preys (FDR ≤ 0.01) were plotted for LMNA-BioID vs LMNA-miniTurbo. The average spectral count value found in controls was subtracted from the detected spectral counts for each prey and the resulting value plotted. Zero values were set to 0.05 to create values suitable for log-transformation of the axes. **b**) LMNA-BioID vs LMNA-turboID. **c)** LMNA-miniTurbo vs LMNA-TurboID.

**Extended Data Figure 11**: Dotplot view of BioID data for mito-ER contact site candidates highlighting recovery of mitochondrial fission machinery, mito-ER tethers and outer mitochondrial membrane proteins. Asterisks on the heat map indicate spectral counts for prey genes corresponding to the bait that were ignored by SAINT as peptides from the bait itself confound accurately evaluating the abundance of itself as an interactor.

**Extended Data Figure 12**: Exploratory analysis of FAM171A1 and MTFR2. BioID was performed on these two baits and the SAINT-processed data was analyzed using the analysis module at humancellmap.org. a) Specificity plot of FAM171A1 showing the high abundance and/or specificity of cytoskeletal proteins. b) Specificity plot of MTFR2 showing the high specificity of proteins involved in mitochondrial dynamics.

**Extended Data Figure 13**: BRD3 relocalization on JQ1 treatment. BirA-tagged BRD3 was treated with vehicle or JQ1 for 24 hours (data from ^42^, and analyzed using the analysis module at humancellmap.org. The Jaccard indices (1 – Jaccard distance) for the top 20 most similar baits were used to create networks in Cytoscape^48^ using an edge-weighted spring-embedded layout. Humancellmap baits are colored based on their expected localization to chromatin or the nucleolus.

**Extended Data Figure 14**: BirA*-FLAG and GFP-BirA*-FLAG control stable cell line, and LMNA-BirA*-FLAG and AIFM1-BirA*-FLAG bait stable cell line immunofluorescence. Negative control cell lines were probed by confocal immunofluorescence (IF) microscopy in HEK293 Flp-In T-REx stable cells to assay for localization of the fusion construct and general biotinylation. Cells were fixed and then probed with an antibody to FLAG epitope and streptavidin for biotinylated proteins (see **Methods** for details). Green channel represents nuclear or mitochondrial staining, red channel for FLAG and the blue channel for streptavidin (biotinylated proteins). Scale bars, 10 μm.

**Supplementary Table 1**: Bait description and bait quality control.

**Supplementary Table 2**: Mass spectrometry samples and results for the core BioID dataset.

**Supplementary Table 3**: Enriched terms for significant proximal proteins.

**Supplementary Table 4**: Previously reported preys.

**Supplementary Table 5:** Factors affecting prey recovery.

**Supplementary Table 6**: Enriched GO cellular component terms in prey-prey heat map clusters.

**Supplementary Table 7**: SAFE localization predictions.

**Supplementary Table 8**: NMF localization predictions.

**Supplementary Table 9**: NMF and SAFE compartment definitions.

**Supplementary Table 10:** Localization prediction metrics.

**Supplementary Table 11**: NMF and SAFE protein domain and motif enrichments.

**Supplementary Table 12**: Benchmarking of NMF and SAFE localization predictions.

**Supplementary Table 13**: Predicted localization validation details.

**Supplementary Table 14:** ER-localized prey exposure to the cytosol and lumen.

**Supplementary Table 15**: Moonlighting and compartment connections.

**Supplementary Table 16**: LMNA profiling with different biotin ligases.

**Supplementary Table 17**: Mass spectrometry samples and results for the ER-Mito BioID dataset.

**Supplementary Table 18**: Mass spectrometry samples and results for the prediction BioID dataset.

**Supplementary Table 19**: BRD3 analysis.

**Supplementary Table 20:** Control sample profiles.

**Supplementary Table 21**: List of reagents.

## Methods

### Selection of compartment markers

We aimed at selecting at least three independent baits (referred to here as “compartment markers”) for all major membrane-bound and membraneless organelles in HEK293 cells, as well as for all cytoskeletal elements. For complex organelles, such as the nucleus and the mitochondrion, distinct markers were selected to profile their major subcompartments (e.g. matrix, inner membrane and outer membrane for the mitochondria). These markers were selected by manual literature curation (e.g. they have previously been used as fluorescent recombinant proteins or sequence tags to mark selected structures), from proteins reported as high quality markers in the Human Protein Atlas^1^, commercially used as compartment markers for immunofluorescence (e.g. Cell Signaling Technology), or following advice from cell biology experts. The list of the constructs used here can be found in **Supplementary Table 1**.

The selection of the BirA*-FLAG location (N-or C-terminus) for each marker was as follows: if the selected marker had previously been used successfully for fluorescent-protein tagging and microscopy, the same tag location was used for BioID. For proteins where such information was not available (e.g. they were used as endogenous markers), the structural organization of the protein was taken into consideration (for example, if a critical domain or motif such as a mitochondrial localization sequence or prenylation motif, was present at one of the termini, the other terminus was used for tagging). For transmembrane domain-containing proteins, membrane topology was analyzed from both the literature and using the Protter tool^49^, and the tag integrated on the side of the membrane where compartment labelling was desired. In six cases, both N and C-terminal fusions of the same protein were generated.

Selected markers were subcloned as in-frame fusions by Gateway cloning in the pcDNA5-FLAG-BirA* backbone^50^ (with fusion of the marker at either the N-or C-terminus). When no appropriate entry Gateway construct was available, entry clones were generated by PCR amplification from cDNA constructs (Mammalian Gene Collection; MGC). “Open” Gateway constructs destined for N-terminal fusions were first “closed” by PCR amplification and recloned as closed entries to prevent cloning scars^51^. Sequence tags^52^ were PCR amplified from relevant cDNA or Gateway ORF clones of the full-length proteins, or from oligo annealing, and inserted into the pcDNA5-FLAG-BirA* backbone. All constructs generated by PCR amplification were validated by Sanger sequencing.

### Cell line generation for BioID

For BioID, the parental cell line, HEK293 Flp-In T-REx 293 (Invitrogen), was grown at 37**°**C in DMEM high glucose supplemented with 5% Fetal Bovine Serum, 5% Cosmic calf serum and 100 U/ml Pen/Strep (growth media). Parental cell lines were routinely monitored for mycoplasma contamination and were authenticated by STR analysis with The Center for Applied Genomics Genetic Analysis Facility (Sick Kids Hospital, Toronto).

For the generation of stable cell lines, HEK293 Flp-In T-REx cells were transfected using the jetPRIME transfection reagent (Polyplus Cat# CA89129-924). Cells were seeded at 250,000 cells/well in a 6-well plate in 2 ml growth media (day 0). The following day (day 1), cells were transfected with 100 ng of pcDNA5-FLAG-BirA* bait construct and 1 μg of pOG44 in 200 μl of jetPRIME buffer mixed with 3 μl of jetPrime reagent (of this mix, 200 μl was added to the cells as per the manufacturer’s protocol). On day 2, transfected cells were passaged to 100 mm plates. On day 3, hygromycin was added to the growth media (final concentration of 200 μg/ml). This selection media was changed every 2–3 days until clear visible colonies were present, at which point the colonies were pooled. Cells were then scaled up to 150 cm^2^ plates. Cells were grown to 70% confluence before induction of protein expression using 1 μg/ml tetracycline, and the media supplemented with 50 μM biotin for protein labelling. Cells were harvested 24 h later as follows: cell media was decanted, cells were washed once with 5 ml PBS, then harvested by scraping in 1 ml PBS. Cells from one or two 150 cm^2^ plates were pelleted at 233 RCF for 5 min, the supernatant aspirated, and pellets frozen on dry ice. Cell pellets were stored at −80°C until further processing.

### BioID

The BirA* enzyme used in this study for profiling compartments was the original BioID enzyme described in ^7^. Two different BioID protocols were implemented and are described below. The protocol used for each bait can be found in **Supplementary Table 2**.

#### Protocol 1 (high stringency washes; highSDS)

Cell pellets from one 150 mm plate were lysed in a modified RIPA buffer containing MgCl2 (modRIPA + MgCl2: 50 mM Tris-HCl pH 7.5, 150 mM NaCl, 1.5 mM MgCl2, 1% Triton X-100, 1 mM EGTA, 0.1% SDS, Sigma-Aldrich protease inhibitors P8340 1:500 (v:v), and 0.5% Sodium deoxycholate) at 1:10 (pellet weight in g: lysis buffer volume in ml). After lysis buffer addition, 1 μl of benzonase (EMD, CA80601-766, 250 U) was added to each sample, and cell pellets were incubated on a nutator at 4°C for 20 min. Lysates were sonicated (3 x 10-second bursts with 2 seconds rest) on ice at 65% amplitude using a Qsonica with a CL-18 probe. Lysates were centrifuged for 30 min at 20,817 RCF at 4°C. After centrifugation, lysate supernatants were added to pre-washed streptavidin-sepharose beads (GE Cat# 17-5113-01; 30 μl bed volume of pre-washed beads per sample), and biotinylated proteins were affinity-purified at 4°C on a nutator for 3 h. After affinity purification, streptavidin sepharose beads were pelleted (400 RCF, 1 min), and the supernatant removed. Streptavidin beads were then transferred to a new microfuge tube in 1 ml of 2% SDS Wash Buffer (2% SDS, 25 mM Tris-HCl pH 7.5). All subsequent washes used 1 ml of the indicated buffer with a centrifugation force of 400 RCF for 1 min. Beads were washed twice with modRIPA +MgCl2 (without protease inhibitors or sodium deoxycholate), and three times with 50 mM ammonium bicarbonate buffer (pH 8). All buffer was removed from the final wash, and 1 μg of mass spectrometry grade trypsin/Lys-C mix (Promega CAT# V5071) in 60 μl of 50 mM ammonium bicarbonate was added to each sample. Proteins were digested on beads overnight at 37 °C on a rotator. The following day, an additional 0.5 μg trypsin/Lys-C mix was added to samples that were further digested at 37 °C on a rotator for 2 h. Each sample was spun down at 400 RCF for 1 min to pellet beads, and the supernatant was transferred to a new 1.5 ml microcentrifuge tube. Beads were then washed with 30 μl of HPLC-grade water (Caledon Laboratory Chemicals CAT# 7732-18-5), centrifuged at 400 RCF for 1 min to pellet beads, and the supernatant pooled with digested peptides collected previously (this step was repeated once). Samples were centrifuged at 16,000 RCF for 5 min and 100 μl was transferred to a new microfuge tube. Samples were acidified by adding 4 μl of 50% formic acid (final concentration of 2% formic acid) and dried in a centrifugal evaporator. Dried peptides were stored at −80 °C.

#### Protocol 2 (lower stringency washes; lowSDS)

This follows the same steps as Protocol 1, except for the details listed below. Cell pellets from two 150 mm plates were lysed in modified RIPA buffer containing EDTA (modRIPA + EDTA: 50 mM Tris-HCl pH 7.5, 150 mM NaCl, 1% Triton X-100, 1 mM EDTA, 1 mM EGTA, 0.1% SDS, Sigma-Aldrich protease inhibitors P8340 1:500 (v:v), and 0.5% Sodium deoxycholate) at 1:10 (pellet weight in g: lysis buffer volume in ml). After affinity purification, streptavidin beads were transferred to a new microfuge tube in 1 ml of modRIPA +EDTA (without protease inhibitors or sodium deoxycholate). All subsequent washes used 1 ml of a buffer with a centrifugation force of 400 RCF for 1 min. The beads were washed once more with modRIPA +EDTA (without protease inhibitors or sodium deoxycholate), twice with an NP-40 wash buffer (10% glycerol, 50 mM HEPES-KOH pH 8.0, 100 mM KCl, 2 mM EDTA, 0.1% NP-40) and three times with 50 mM ammonium bicarbonate (pH 8) buffer. All of the buffer was removed from the final wash, and 1 μg of mass spectrometry grade trypsin (Sigma-Aldrich T6567) in 200 μl of 50 mM ammonium bicarbonate was added to each sample. Samples were digested on beads overnight at 37 °C on a rotator. After the addition of an additional 0.5 μg of trypsin and 2 h incubation, the digested peptides were transferred to a new 1.5 ml microcentrifuge tube. Beads were then washed with 150 μl of HPLC-grade water (Caledon Laboratory Chemicals CAT# 7732-18-5), centrifuged at 400 RCF for 1 min to pellet beads, and the supernatant pooled with digested peptides collected previously. The water wash and collection of the supernatant were repeated once more. Digested peptides were centrifuged at 16,000 RCF for 5 min and 470 μl collected into a new microfuge tube. Samples were dried in a centrifugal evaporator, and dried peptides were stored at −80 °C.

### Mass spectrometry analysis

Dried peptides were resuspended in 20 μl of 5% formic acid and centrifuged at 16,000 RCF for 1 min. 5 μl were injected via autosampler in a 12 cm analytical fused silica capillary column (0.75 μm internal diameter, 350 μm outer diameter). The column was made in house using a laser puller (Sutter Instrument Co., model P-2000; heat = 280, FIL = 0, VEL = 30, DEL = 200), packed with C18 reversed-phase material (Reprosil-Pur 120 C18-AQ, 3 μm; Dr. Maische), and connected in-line to a NanoLC-Ultra 2D plus HPLC system (Eksigent, Dublin, USA). The system was equipped with a nanoelectrospray ion source (Proxeon Biosystems, Thermo Fisher Scientific) delivering the sample to an Orbitrap Elite Hybrid Ion Trap-Orbitrap mass spectrometer (Thermo Fisher Scientific). The HPLC program delivered the following percentages of buffer B (0.1% formic acid in acetonitrile) to buffer A (0.1% formic acid in water) at the described flow rates over a 130 min gradient. The start of the HPLC program loaded the sample onto the column with a flow rate of 400 μl/min with 5% buffer B for 14 min followed by a drop in flow rates from 400 μl/min to 200 μl/min using a linear gradient from 5% to 2% buffer B for 1 min. Next, a linear gradient from 2% to 35% buffer B began eluting the sample into the mass spectrometer at 200 μl/min for 90 min, followed by another linear gradient from 35 to 80% buffer B over 5 min, and maintaining 80% buffer B for 5 min to elute the remaining analytes. The final stages of the HPLC program had a flow rate of 200 μl/min using a linear gradient from 80% to 2% buffer B over 3 min, and a quick re-equilibration of the column for 12 min at 200 μl/min with 2% buffer B.

The Orbitrap Elite Hybrid Ion Trap-Orbitrap mass spectrometer was operated with Xcalibur 2.0 software in data-dependent acquisition mode with the following parameters: one centroid MS (mass range 400 to 2000) followed by MS2 on the top 10 most abundant ions with a dynamic exclusion of 20 s (general parameters: activation type = CID, isolation width = 2 m/z, normalized collision energy = 35, activation Q = 0.25, activation time = 10 ms. The minimum signal required was 1000, the repeat count = 1, repeat duration = 30 s, exclusion size list = 500, exclusion duration = 15 s, exclusion mass width (Da) = low 0.6, high 1.2). To decrease carry over between samples on the autosampler, the analytical column was washed three times using a “sawtooth” gradient of 35% acetonitrile with 0.1% formic acid to 80% acetonitrile with 0.1% formic acid, holding each gradient for 5 min, three times per gradient. Following washes, quality control on the column and machine performance were assessed by loading 30 fmol BSA tryptic peptide standard (Michrom Bioresources Inc. Fremont, CA) with 60 fmol α-Casein tryptic digest. The HPLC program for the quality control ran a shortened 60 min gradient with the following percentages of buffer B and flow rates: 9 min at 400 μl/min with 5% buffer B, 1 min going from 400 μl/min to 200 μl/min using a linear gradient from 5 to 2% buffer B, 30 min at 200 μl/min using a linear gradient from 2 to 35% buffer B, 5 min at 200 μl/min using a linear gradient from 35 to 80% buffer B, 5 min at 200 μl/min with 80% buffer B, 5 min at 200 μl/min using a linear gradient from 80 to 2% buffer B and 5 min at 200 μl/min with 2% buffer B.

### Mass spectrometry data analysis

Mass spectrometer raw files were converted to mzML using ProteoWizard (3.0.4468)^53^ and analyzed using the iProphet^54^ pipeline implemented within ProHits^55^ as follows. The database consisted of the HEK293 sequences in the RefSeq protein database (version 57) supplemented with “common contaminants” from the Max Planck Institute http://141.61.102.106:8080/share.cgi?ssid=0f2gfuB and the Global Proteome Machine (GPM; http://www.thegpm.org/crap/index.html) with the addition of sequences from common fusion proteins and epitope tags. The search database consisted of forward and reverse sequences (labeled “gi|9999” or “DECOY”); in total, 72,226 entries (including decoys) were searched. Spectra were analyzed separately using Mascot (2.3.02; Matrix Science) and Comet (2012.01 rev.3)^56^ for trypsin specificity with up to two missed cleavages; deamidation (NQ) or oxidation (M) as variable modifications; single-, double-, and triple-charged ions allowed, mass tolerance of the parent ion to 12 ppm; and the fragment bin tolerance at 0.6 amu. The resulting Comet and Mascot search results were individually processed by PeptideProphet^57^, and peptides were assembled into proteins using parsimony rules first described in ProteinProphet^58^ into a final iProphet protein output using the Trans-Proteomic Pipeline (TPP; Linux version, v0.0 Development trunk rev 0, Build 201303061711). TPP options were 1) general options: −p0.05 −x20 −PPM −d”DECOY”, 2) iProphet options:–ipPRIME and 3) PeptideProphet options: –pP. All proteins with a minimal iProphet protein probability of 0.05 were parsed to the relational module of ProHits. Note that for analysis with SAINT (see below), only proteins with iProphet protein probability ≥ 0.95 were considered, corresponding to an estimated protein level false-discovery rate (FDR) of approximately 0.5%.

### SAINT file processing

For each prey protein identified in an affinity purification experiment, SAINT calculates the probability of it being a true interaction by using spectral counting (semi-supervised clustering, using a number of negative control runs). SAINTexpress^14^ analysis was performed using version exp3.6.1 with two biological replicates per bait. Two separate SAINT analyses were performed for the two BioID protocols. For the baits used with BioID Protocol 1, 322 bait protein samples (162 baits) were analyzed alongside 70 negative control runs, consisting of purifications from untransfected cells or cells expressing BirA*-FLAG or BirA*-FLAG-GFP. For BioID Protocol 2, 52 bait protein samples (26 baits) were analyzed alongside 16 negative control runs, consisting of purifications from untransfected cells or cells expressing BirA*-FLAG or BirA*-FLAG-GFP. No compression of the controls was performed and default parameters for SAINTexpress were used. A 1% Bayesian FDR cut-off was used to select confident proximity interactors relative to the expected spectral count distribution seen in control samples. All prey proteins detected in controls samples and enriched GO terms for the top preys in these samples can be found in **Supplementary Table 20**. The two SAINT files for the core dataset were combined into a single file for downstream analysis, and non-human contaminants were removed from the final report, as were baits with less than 5 significant preys. SAINTexpress was also used in a separate analysis for the proximity proteomes of the “Prediction” baits; Protocol 1 controls described above were used for this analysis, using the same parameters as above.

### Immunofluorescence (IF) microscopy for bait quality control

For quality control of stable cell lines expressing BirA*-FLAG-tagged baits, HEK293 Flp-In T-REx cells were seeded directly on 12 mm poly-L-lysine coated coverslips (Corning, Product # 354085). The next day, cells were treated with 1 μg/ml tetracycline and media was supplemented with 50 μM biotin for 24 h. Media was aspirated, and cells were washed with PBS supplemented with 200 μM CaCl2, 100 μM MgCl2, prior to fixation with 4% formaldehyde in PBS for 10 min, and washing three times in TBS-T (Tris-buffered saline and 0.1% v/v). The cells were then treated for 10 min in permeabilization buffer (0.1% Triton X-100 in TBS-T), followed by 3 washes in TBS-T and incubation at room temperature in blocking buffer (5% BSA w/v in TBS-T). Samples were incubated with primary antibodies in blocking buffer in a humidified chamber for 1 h: anti-FLAG M2 (1:2000 dilution, Sigma Aldrich, F3165) and an endogenous compartment marker antibody from rabbit (**Supplementary Table 21** for list of antibodies used), or anti-FLAG from rabbit (1:500 dilution, Sigma Aldrich, F7425) and an endogenous compartment marker antibody from mouse. All samples were then washed 3 times in blocking buffer before incubation with blocking buffer containing secondary antibodies in a dark, humidified chamber for 1 h with one of the combination of antibodies and dyes listed here: (1) anti-rabbit coupled to Alexa Fluor 488 (1:1000, Invitrogen, A11034), anti-mouse coupled to Alexa Fluor 555 (1:1000; Invitrogen, A21422), Streptavidin-coupled to Alexa Fluor 647 (1:2500, Invitrogen, S32357); (2) anti-mouse coupled to Alexa Fluor 488 (1:1000, Invitrogen, A11001), anti-rabbit coupled to Alexa Fluor 555 (1:1000; Invitrogen, A21428), Streptavidin-coupled to Alexa Fluor647 (1:2500, Invitrogen, S32357); (3) anti-mouse coupled to Alexa Fluor 555 (1:1000; Invitrogen, A21422), DAPI (1:2000), Streptavidin-coupled to Alexa Fluor 647 (1:2500, Invitrogen, S32357); or (4) anti-mouse coupled to Alexa Fluor555 (1:1000, Invitrogen, A21422), Phalloidin-coupled to Alexa Fluor 488 (1:1000, Invitrogen, A12379), Streptavidin-coupled to Alexa Fluor 647 (1:2500, Invitrogen, S32357). Following incubation, samples were washed 3 times with TBS-T. Each coverslip was mounted on a glass slide using ~4 μl of ProLong Gold Antifade Mountant (Thermo Fisher Scientific, CAT #P36930). Samples were then cured, lying flat, overnight in the dark, followed by storage in the dark at 4°C. Images were acquired on a Nikon C1Si Confocal Microscope using a 60x objective lens magnification and 3x field zoom.

In some instances, ice-cold methanol (MeOH) was used as a fixative to better visualize microtubules and facilitate the use of specific antibodies only amenable to MeOH fixation conditions. Ice-cold MeOH addition and incubation at −20°C for 30 min was used to fix and permeabilize cells after the first initial wash. After the cells were washed 3 times with TBS-T, the protocol continued as described above with the addition of blocking buffer. When Wheat Germ Agglutinin (WGA)-coupled to Alexa Fluor488 (1:250, Invitrogen, W11261) was used as a counterstain, all steps were performed with samples chilled on ice, employing ice-cold buffers and in the dark. After the initial wash, cells were incubated with a solution containing WGA-coupled to Alexa Fluor488 in PBS containing 200 μM CaCl2, 100 μM MgCl2 for 10 min. After the samples were washed twice with this solution, the protocol was as described for formaldehyde fixation.

The localization of negative controls (BirA*-FLAG and GFP-BirA*-FLAG) can be found in **Extended Data Figure 14.**

### GO enrichment analysis

GO enrichments were performed using g:Profiler^59^. Enrichments were performed considering gene lists as unordered, allowing only genes with annotations, using all significant proximal interactors as background, a max p-value of 0.01 and the g:SCS multiple test correction method. For bait QC, it should be noted that DHFR2 did not match its expected compartment enrichment but was allowed into our analysis pipeline as its large list of proximal interactors was deemed to be informative for localization purposes. NPM1 and KDM1A had an expected GO:CC enrichment profile when the max p-value was relaxed from 0.01 to 0.05.

### Databases used for analysis

The BioGRID^16^ human database v3.5.169 was downloaded on 13/2/2019. Human gene annotations were downloaded from Gene Ontology (GO) on 15/2/2019 (GO version date 1/2/2019). The GO hierarchy (release date 13/2/2019) was downloaded from GO^60,61^ on 15/2/2019. The UniProt database^62^ was downloaded on 21/2/2019. The IntAct^17^ human database was downloaded on 13/2/2019. Human protein domain annotations and motifs were retrieved from Pfam^63^ (version 32) on 21/2/2019. ProteomicsDB^19^ was queried for protein expression information on 1/14/2020. Text mining data was downloaded from the Compartments database^64^ on 1/21/20.

### Jaccard index

The Jaccard index is the overlap between two sets (*A, B*) calculated as

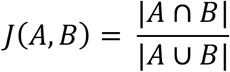

The Jaccard distance is defined as 1 – *J*(*A,B*).

### Control-subtracted length-adjusted spectral counts (CLSC)

The prey proximity order for each bait was determined from the prey’s control-subtracted length-adjusted spectral counts. For bait *i* and prey *j* this value was calculated by first subtracting the prey’s average spectral count found in control samples from its abundance with bait *i,* then multiplying by the median prey length of all preys across bait *i* and dividing by the length of prey *j*.

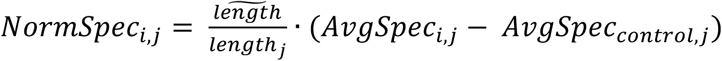

### Prey specificity

The specificity of a prey for a particular bait at the humancellmap is calculated as the spectral count detected with the bait, divided by the mean spectral count across all other baits or against the top ten most similar baits as indicated on the analysis report. Whenever available, the average spectral count detected in control samples is subtracted from the prey count prior to calculating the specificity, as was done for all specificity metrics reported in this study.

### Prey-prey correlation

The SAINTexpress file was processed using the correlation tool at ProHits-viz^46^ with an FDR score filter of 0.01 and an abundance cut-off of 0. If a prey passed the FDR cut-off for one bait, its abundance across all other baits was used in the analysis. Control average values were subtracted from replicate spectral counts and these control-subtracted values used for correlation. After Pearson correlation scores were calculated between preys, complete-linkage clustering was performed using the Euclidean distance between preys, and cluster order was optimized using the CBA package in R.

### SAFE

A network was built from prey-prey correlation data using ProHits-viz as described above. Networks were built in Cytoscape^48^, version 3.6.1 using a spring embedded layout. All preys passing an FDR cutoff of 0.01 were included in this analysis. After performing correlation, we considered preys to be interaction pairs if they passed a required correlation cut-off. This cutoff was set to 0.5 to 0.9 in increments of 0.05 for testing with SAFE^65^ as we could not know a priori what an ideal cut off would be although manual assessment suggested something in this range would be suitable. SAFE requires annotations for network nodes and for each node we created a list of all known GO cellular compartment terms supplemented with their parent terms. When running SAFE, we also tested several percentile neighbourhood radii for each network, ranging from 3 to 10 in increments of 0.5. With these parameters, we sought to maximize the number of preys being assigned to a domain with a known GO term for that prey. A prey was considered assigned to a correct domain if one of its GO terms (or a parent of those terms) was found within the terms assigned to its predominant domain. After manually inspecting the SAFE results, we felt the optimal annotation was generated from the network built with a correlation cut-off of 0.65 with a neighbourhood radius of 4.5. This resulted in a network with 24 domains (one of which is “unknown”), where 60.2% (2351/3903) of genes were assigned to a domain with a known GO term. The complete definition of each domain was determined by the GO cellular compartment terms resulting from an enrichment of all preys with a primary localization to the domain in question. We also selected a representative term(s) for each domain as its compartment ID for localization and assessment purposes.

### NMF

Non-negative matrix factorization (NMF) is an approach to create a compressed and simplified version of an *n* × *m* dimension matrix **V**, such that **V** ≈ **WH**, where **W** has dimensions *n* × *r* and **V** has dimensions *r* × *m*, and both matrices consist entirely of non-negative entries^66^. Given an interaction matrix of *n* preys and *m* baits, where *V_i,j_* is the spectral count of prey *i* with bait *j,* the minimal rank *r* of the factorization is sought that sufficiently summarizes this input matrix. In our case, we seek r << m. The matrix **W** can then be thought of as a compressed form of our input matrix, whereby instead of displaying a prey’s profile across all baits, it shows how preys profile across ranks. A simple way to think of a rank in the context of our dataset is that it may represent a collection of baits that convey redundant information. In contrast to the input matrix that may show several data points indicating a prey is detected highly with each nuclear bait, for example, we might expect a single entry in the matrix indicating it was detected highly in the nucleus. Preys that behave similarly across baits would be expected to have similar profiles across ranks. Preys that only behave similarly across a subset of baits would still be expected to show a similar profile across a single or subset or ranks, while being free to show a different profile across the remaining ranks. Our input matrix had dimensions 4424 *×* 192 for the 192 baits in the data set and 4424 preys passing an FDR cutoff of 1%. Prey spectral counts had their average value in controls subtracted and were then rescaled from 0–1 across baits as we wanted each prey to be considered of equal weight. NMF, as implemented optimizing the squared Frobenius norm initialized by Nonnegative Double Singular Value Decomposition (NNDSVD) with L1 regularization in the *-scikit-learn* Python package^67^, version 0.18.1, was then performed on this matrix for *r* = 10, 11…30. For each NMF run, GO cellular compartment terms were assigned to the resulting ranks by taking the top preys for each rank in the **W** matrix (up to 100 maximum) and profiling with g:Profiler^68^ using our complete prey list as background. A prey could contribute to the enrichment process in an NMF rank if it was most abundant in that rank or within 25% of its maximum within that rank, and if it had a value of at least 0.25. These values were set to try and ensure there was sufficient evidence that a prey truly belonged to a rank. To determine the optimal number of ranks to use for NMF, we sought to maximize the number of preys assigned a known localization and minimize the overlap in GO terms between ranks. A prey was considered assigned to a correct rank if one of its known GO terms (or a parent of those terms) was found within the terms assigned to its rank. To determine the overlap in GO terms between ranks, we calculated the Jaccard distance between GO terms for each pair of ranks (where 0 would indicate complete overlap and 1 no overlap). While several NMF ranks performed well, we selected 20 ranks after manual inspection. Analysis with 20 ranks resulted in 87.6% of preys assigned to a rank with a previously known GO term and 74.8% of preys assigned to a rank where one of the top 5 GO terms was previously known, with the worst rank overlap at a Jaccard distance of 0.31. After defining the optimal rank number, each prey was assigned to its best rank for visualization and assessment purposes, and a representative GO term or terms was/were chosen to identify the rank, and also for visualization and assessment purposes. Since at most only the top 100 preys in a rank were used for its definition, we used the remaining preys localized to the rank to assess the ability of this approach to correctly localize proteins. 48.0% of these preys were localized to a previously known compartment (based on GO:CC annotations, **Supplementary Table 8**), giving us confidence in the procedure.

A network was built from the pairwise prey Euclidean distance matrix derived from the NMF **W** matrix using t-Distributed Stochastic Neighbor Embedding (t-SNE)^45^. t-SNE was performed using the Matlab script available at http://lvdmaaten.github.io/tsne/. It was run with the number of initial dimensions equal to the number of NMF ranks (20) and a perplexity of 20 for a maximum of 1000 iterations.

### Information content

The information content (IC) of each GO cellular component term was calculated as −log(*p*), where *p* is the probability a gene has an annotation, i.e. the number of genes with the annotation divided by the total number of genes in GO. Annotations occurring in 1% of genes or less (189 / 18858 total genes in GO) were placed in our highest specificity IC tier (bin 1). Bins 2-5 corresponded to annotations occurring in 2%, 10%, 25% or > 25% of genes.

### Dataset comparison

The Human Protein Atlas (HPA) subcellular localization data was downloaded on 15/3/2019 from www.proteinatlas.org/about/download, and is based on the Human Protein Atlas^69^ version 18.1 and Ensembl version 88.38. All HPA entries in the subcellular localization table have an associated gene name and all localization terms are based on GO. Fractionation-based localizations from Christoforou et al.^2^ were retrieved from Supplementary dataset 1, tab 2, column AI (“Final Localization Assignment”). Their localization terms were mapped to the closest GO term. Although their dataset is for mouse genes, more localizations were known if we assumed their genes were human and compared against the human GO database, so this was used for our assessment. Fractionation-based localizations from Itzhak et al.^3^ were retrieved from Supplementary file 1, tab 3, columns H, K and N (“Compartment Prediction”, “Sub compartment prediction” and “Global classifier”). Columns H and K were first merged. Missing predictions, “No Prediction” and “Large Protein Complex” assignments were ignored and instead the classification from column N was used. Localization terms were mapped to the closest GO term. Predictions from canonical isoforms were used when possible or alternative isoforms if the canonical isoform had no prediction. In the case of only non-canonical isoforms, the most specifically localized isoform was used. All genes from these datasets with their assigned and corresponding GO IDs are listed in **Supplementary Table 12**. Localization tiers were defined using the information content of each GO term as defined in the “Information Content” section. When genes were assigned multiple localizations, the lowest information content term (i.e. least specificity) was used for binning that gene into a localization tier.

### Enrichments

Enrichment scores (p-values) for domain and motif enrichment were calculated for each NMF rank and SAFE domain using Fisher’s exact test. Of the 4424 genes in our NMF analysis, 4368 had domain information available and 4301 had motif information available in Pfam. Of the 3903 genes in our SAFE analysis, 3855 had domain information available and 3809 had motif information available in Pfam. All genes with available information were used as background for the enrichment tests. The FDR was controlled by using the Benjamini–Hochberg procedure for an FDR of 1%.

### Validation of localization predictions by immunofluorescence microscopy and BioID

For prediction validation we prioritized proteins without a clear annotation, for example proteins annotated as an Open Reading Frame, ORF, or simply as “family with sequence similarity”, FAM. We also focused on families that share domains or structural features for which multiple members were present on our map, but with different predicted localization. These included proteins annotated as solute carriers (SLC), transmembrane proteins (TMEM), and proteins that contain a Rab small GTPase domain. We only selected proteins where the NMF and SAFE predictions were in agreement and for which we could readily access a fulllength cDNA or ORF clone locally. Selected targets for validation were cloned in Gateway compatible pcDNA5-GFP and pcDNA5-FLAG-BirA* backbones (with tags at either N-or C-terminus as described for the selection of bait quality control above) and localizations validated by immunofluorescence microscopy and GO enrichment as described above (see **Supplementary Table 13** for the list of tested baits).

GFP-tagged constructs were transiently transfected into HeLa cells (ATCC, CCL-2) using the jetPRIME transfection reagent (Polyplus Cat# CA89129-924). Cells were seeded at 250,000 cells/well in a 6-well plate in 2 ml growth media. The following day, cells were transfected with 400 ng of pcDNA5-GFP-tagged construct and 40 μl of jetPRIME buffer mixed with 0.8 μl of jetPrime reagent. The next day formaldehyde fixation, as described above, was used with the following alterations. Samples were incubated with primary antibodies in blocking buffer in a humidified chamber for 1 h. The primary antibodies used were anti-GFP from mouse (1:500 dilution, Roche, CAT# 11814460001) and an endogenous compartment marker antibody from rabbit (refer to **Supplementary Table 21** for list of antibodies used), or anti-GFP from rabbit (1:2000 dilution, abcam, ab290) and an endogenous compartment marker antibody from mouse. Samples were then incubated with blocking buffer containing secondary antibodies in a dark, humidified chamber for 1 h with one of the combination of antibodies and dyes listed here: (1) DAPI (1:2000), anti-rabbit coupled to Alexa Fluor 488 (1:1000, Invitrogen, A11034), anti-mouse coupled to Alexa Fluor 555 (1:1000; Invitrogen, A21422); (2) DAPI (1:2000), antimouse coupled to Alexa Fluor 488 (1:1000, Invitrogen, A11001), anti-rabbit coupled to Alexa Fluor 555 (1:1000; Invitrogen, A21428); or (3) DAPI (1:2000), anti-rabbit coupled to Alexa Fluor 488 (1:1000, Invitrogen, A11034), Phalloidin-coupled to Alexa Fluor647 (1:1000, Invitrogen, A22287). Images were acquired on a Nikon C1Si Confocal Microscope using a 60x objective lens magnification and 1x or 2x field zoom.

BioID was performed on selected targets as described above for cell line generation, BioID Protocol 1, mass spectrometry data analysis and SAINT file processing. For the baits used with BioID Protocol 1, 20 bait protein samples (10 baits) were analyzed alongside 74 negative control runs, consisting of purifications from untransfected cells or cells expressing BirA*-FLAG, or BirA*-FLAG-GFP. GO enrichments were performed using g:Profiler^59^. Enrichments were performed considering gene lists as unordered, allowing only genes with annotations, using a max p-value of 0.05 and the g:SCS multiple test correction method.

Confidence levels of co-localization immunofluorescence images with respect to predicted localizations were assessed and corroborated by three individuals. Confidence rankings were annotated as follows: “supported primary” indicates proteins that matched the primary NMF and SAFE prediction; “supported consistent” indicates proteins that matched the primary NMF and SAFE prediction but did not have an endogenous compartment marker for the immunofluorescence microscopy; “contradiction” indicates proteins that failed to localize to the predicted localizations made by NMF and SAFE; “inconclusive” indicates proteins that had no clear subcellular compartment localization.

### Cell culture for mitochondrial fragmentation assays

Primary fibroblasts (Cell bank at Montreal Children’s Hospital) and HeLa cells were grown in high-glucose DMEM supplemented with 10% fetal bovine serum, at 37°C in an atmosphere of 5% CO2. Stealth RNAi duplex constructs (Invitrogen) were used for transient knockdown of C18orf32 and CHMP7 in primary fibroblasts or HeLa cells. Stealth siRNA duplexes at 12 nM were transiently transfected into cells using Lipofectamine RNAiMAX (Invitrogen 13778-150), according to the manufacturer’s specifications. The transfection was repeated on day 3 and the cells were imaged for mitochondrial morphology analysis on day 6.

### Mitochondrial fragmentation assays

For IF experiments for assaying mitochondrial fragmentation, candidate proteins were GFP-tagged and the constructs were transiently transfected into HeLa cells. HeLa cells were transfected using the jetPRIME transfection reagent (Polyplus Cat# CA89129-924). Cells were seeded at 250,000 cells/well in a 6-well plate in 2 ml growth media. The following day, cells were transfected with 400 ng of pcDNA5-GFP-tagged construct and 40 μl of jetPRIME buffer mixed with 0.8 μl of jetPrime reagent. The next day an IF protocol with FA fixation, described above, was used with the following alterations. Samples were incubated with primary antibodies in blocking buffer in a humidified chamber for 1 h. The primary antibodies used were anti-GFP from mouse (1:500 dilution, Roche, CAT# 11814460001) and anti-COXIV from rabbit (1:250, Cell Signaling Technology, Product# 4850). Samples were then incubated with blocking buffer containing secondary antibodies in a dark, humidified chamber for 1 h with anti-mouse coupled to Alexa Fluor 488 (1:1000, Invitrogen, A11001), anti-rabbit coupled to Alexa Fluor 555 (1:1000; Invitrogen, A21428) and Concanavalin A coupled to Alexa Fluor 647 (1:200, Invitrogen, C21421). Images were acquired on a Nikon C1Si Confocal Microscope using a 60x objective lens magnification. Experiments and image acquisition were separate independent experiments done in triplicate, with an average of n=149 cells per GFP-tagged protein. Mitochondrial fragmentation was quantified manually as deviations from WT mitochondrial staining compared to controls (HeLa cells untransfected or with GFP-alone). Statistical confidence of mitochondrial fragmentation was calculated using the Student’s t-test.

Primary fibroblasts were fixed in warm 4% formaldehyde (FA) in PBS at room temperature for 20 min, then washed three times with PBS before cells were permeabilized in 0.1% Triton X-100 in PBS, followed by three washes in PBS. The cells were then blocked with 3% bovine serum albumin (BSA) in PBS, followed by incubation with primary antibodies (rat anti-KDEL and mouse anti-Cytochrome C, refer to **Supplementary Table 21**) in 3% BSA in PBS for 1 hr at room temperature. After three washes with 3% BSA in PBS, cells were incubated with the appropriate anti-species secondary antibodies coupled to Alexa fluorochromes (1:2000, Invitrogen, **Supplementary Table 21**) for 30 min at room temperature. After three washes in PBS, coverslips were mounted onto slides using fluorescence mounting medium (Agilent Dako). Stained cells were imaged using a 100x objective lenses (NA1.4) on an Olympus IX81 inverted microscope with appropriate lasers using an Andor/Yokogawa spinning disk system (CSU-X), with a sCMOS camera. Mitochondrial network morphology was manually classified, in a blinded manner, as fused, intermediate, or fragmented. For every knockdown condition and controls, 100 −150 cells were analyzed, and experiments were done three times independently. Error bars represent mean ± standard deviation.

### BioID, mass spectrometry analysis and SAINT file processing for mitochondria-ER contact sites

BioID was performed on selected mito-ER candidates as described above for cell line generation, BioID Protocol 1, mass spectrometry data analysis and SAINT file processing. For the baits used with BioID Protocol 1, 20 bait protein samples (10 baits) were analyzed alongside 74 negative control runs, consisting of purifications from untransfected cells or cells expressing BirA*-FLAG, or BirA*-FLAG-GFP. GO enrichments were performed using g:Profiler^59^. Enrichments were performed considering gene lists as unordered, allowing only genes with annotations, using a max p-value of 0.05 and the g:SCS multiple test correction method.

