## Supplementary material for "A proximity-dependent biotinylation map of a human cell: an interactive web resource": Website help documentation

### HELP PAGES

The Human Cell Map is a BioID based map of the human HEK 293 cell. Its aim is to provide a sub-compartment resolution view of the cell. At the cell map you can explore our results or upload your own dataset to compare against ours.

Use the navigation menu at the left to find the help you're looking for. Any questions not answered by this guide should be sent to.

Note for mobile users. This help documentation describes the features available on the desktop version of this site. The mobile version that you are viewing is limited to searching and viewing gene reports.

### DATASET

Here you can find summary statistics as well as databases that were used for generating the cell map.

#### VERSION 1 (CURRENT)

##### Databases

The database used for searching MS files contained the human and adenovirus complements of the RefSeq protein database (version 57) supplemented with common contaminants from the Max Planck Institute and the Global Proteome Machine as well as sequences from common fusion proteins and epitope tags:

- BirA-R118G (UniProt: H0QFJ5)
- Streptavidin (UniProt: P22629)
- GST26 (UniProt: P08515)
- mCherry (UniProt: V9VHH0)
- GFP (UniProt: P42212)
- Lysyl endopeptidase (UniProt: Q9HWK6)

The sequence database consisted of forward and reversed sequences. In total 72, 226 sequences were searched.

##### Summary Statistics

- baits profiled: 192
- unique bait genes: 180
- total interactions: 36038
- unique interactions: 35902
- unique preys: 4424

### EXPLORE

The exploration section of the cell map provides access to our data. You can explore 2D representations of our NMF and SAFE results from the **Map** tab. The **Browse** tab allows you to explore results for specific organelles. You can also search for specific genes from the **Search** tab.

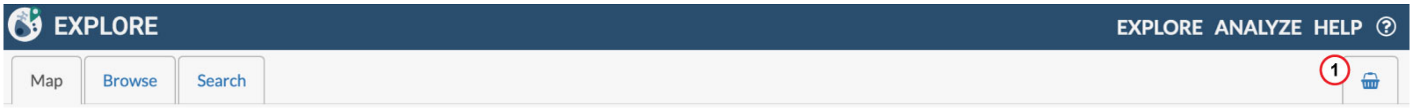


**Explore data at the cell map by clicking on one of the available tabs**

1 Analysis basket for visualizing baits selected while exploring

### EXPLORE

#### MAP/NETWORK

A 2D representation of the cell map can be viewed on the **Map** tab of the Explore section. These interactive maps were generated using NMF and SAFE (please see our manuscript for details). In brief, NMF was used to compress our bait-prey matrix into an organelle-prey matrix and this was converted to a 2D map using t-SNE. For SAFE, proximal interactions were determined via prey-prey Pearson correlation of spectral count values across baits, where preys with a correlation score above a specified cut-off were classified as a proximal interaction pair. This list of interactions was converted to a network using Cytoscape with a spring embedded layout and this network was annotated by SAFE.

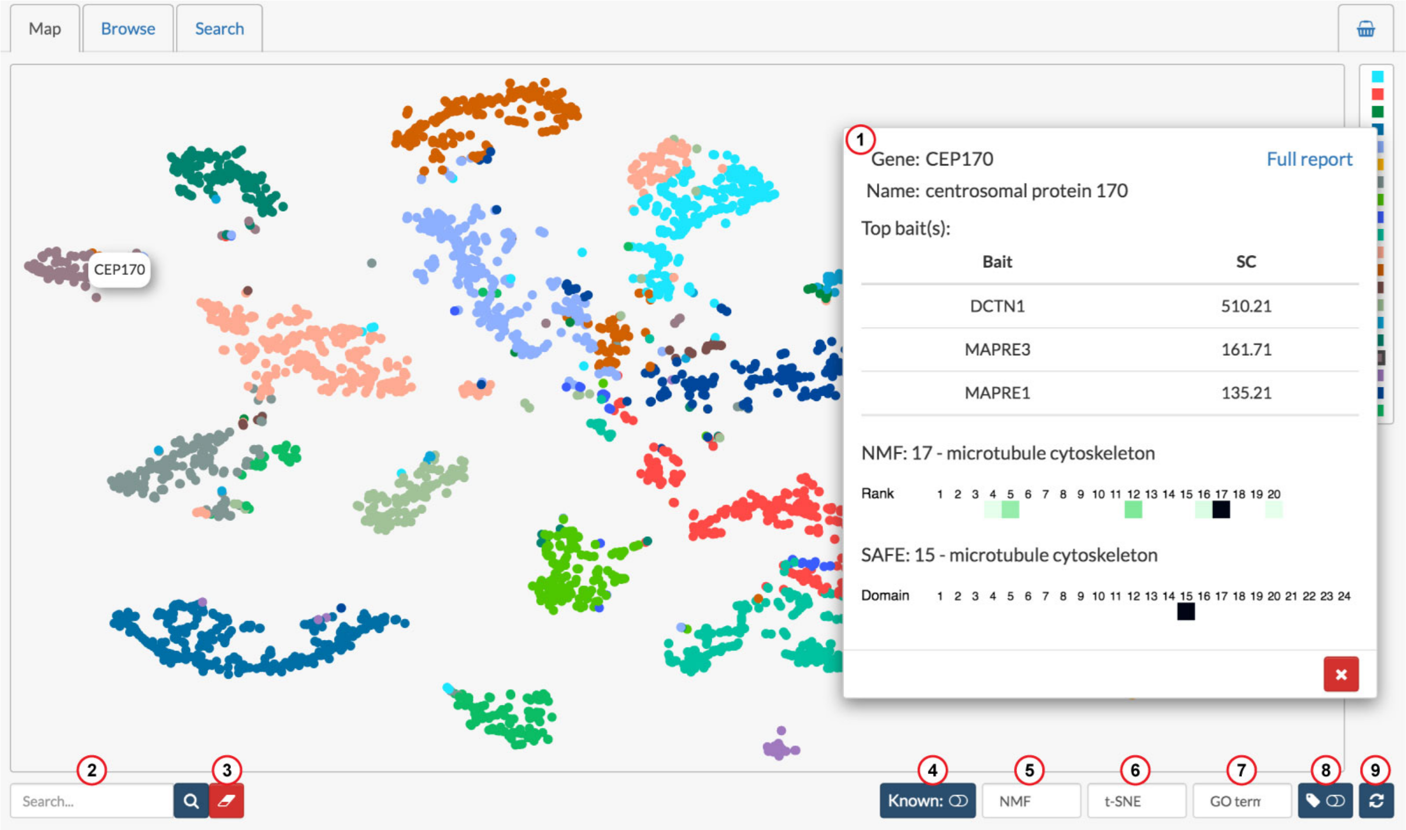


**t-SNE NMF cell map displaying a mini report for CEP170**

1. Mini report for a gene
2. Search for genes on the map
3. Erase gene labels
4. Toggle on/off nodes with previously known localization
5. Switch between NMF and SAFE map
6. Display t-SNE or correlation-based network
7. Change the term type shown when hovering over the legend (GO-term, domains or motifs)
8. Display all node labels

(9) Reset network view

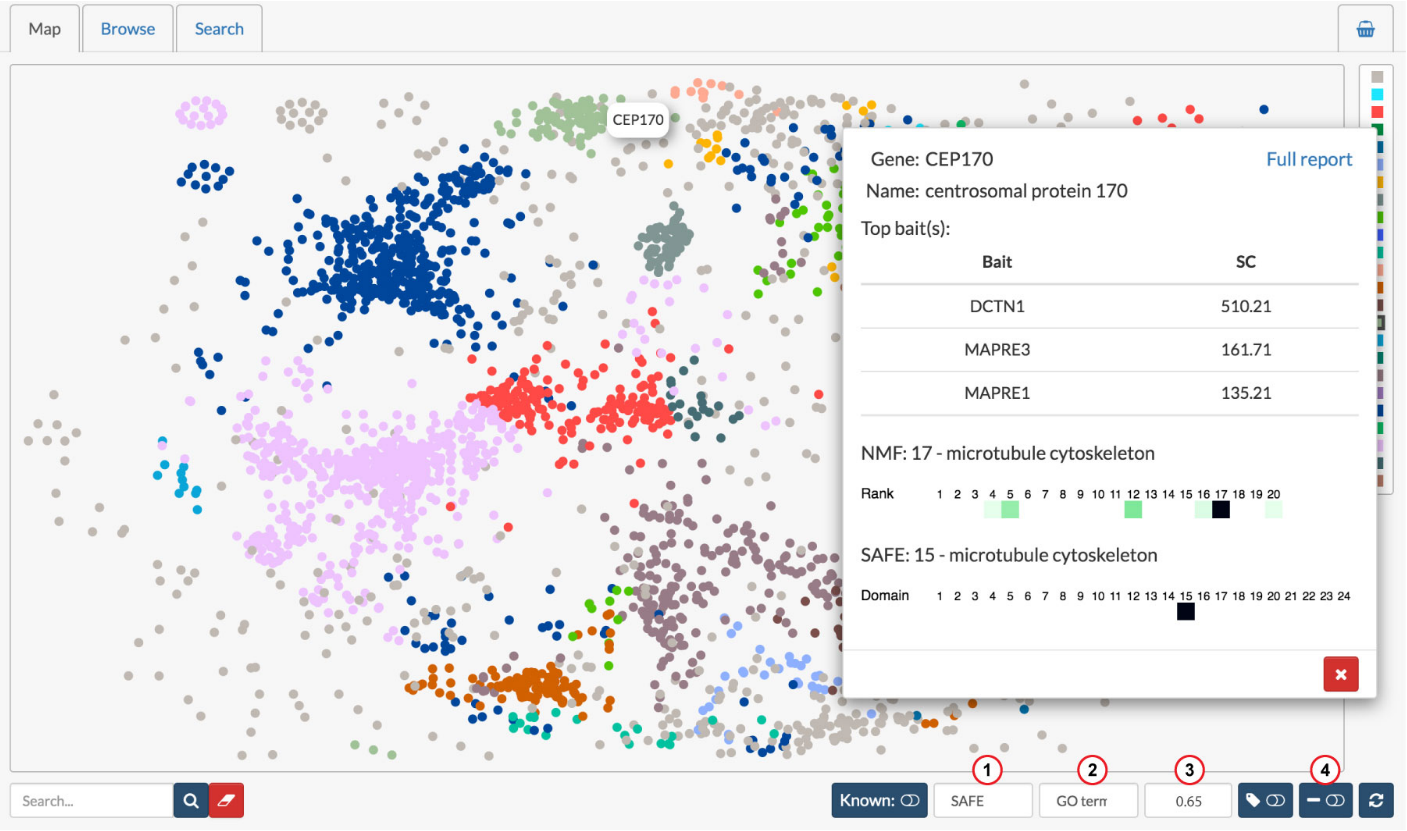


**SAFE cell map displaying a mini report for CEP170**

(1) Switch between NMF and SAFE map

(2) Change the term type shown when hovering over the legend (GO-term, domains or motifs)

(3) Change the correlation cut-off used for defining an interaction pair. A higher cut-off can help to resolve protein complexes

(4) Toggle network edges

Node colour on the map indicates the assigned localization. Hovering over a node will highlight its localization in the legend and will also indicate if the localization was previously known (the node colour will switch to black if not previously known). Hovering over a cell on the legend will display a tooltip with enriched GO terms associated with this localization, with the primary localization shown in bold. GO terms are sorted from most to least enriched on this list.

Clicking on a node will open a mini report about the gene. Control-clicking (Command ⌘ for Mac users) on a node will label the node with its name. Clicking on a cell in the legend will open a full report for that organelle. You can zoom using the mouse wheel and pan around the image by holding the left mouse button or the scroll wheel.

#### References:


- Cytoscape - PMID:14597658
- NMF - PMID:10548103
- SAFE - PMID:27237738
- t-SNE

### EXPLORE

#### BROWSE

From the **Browse** tab you can view the organelles that were profiled and use the cartoon for navigating to organelle-specific reports.

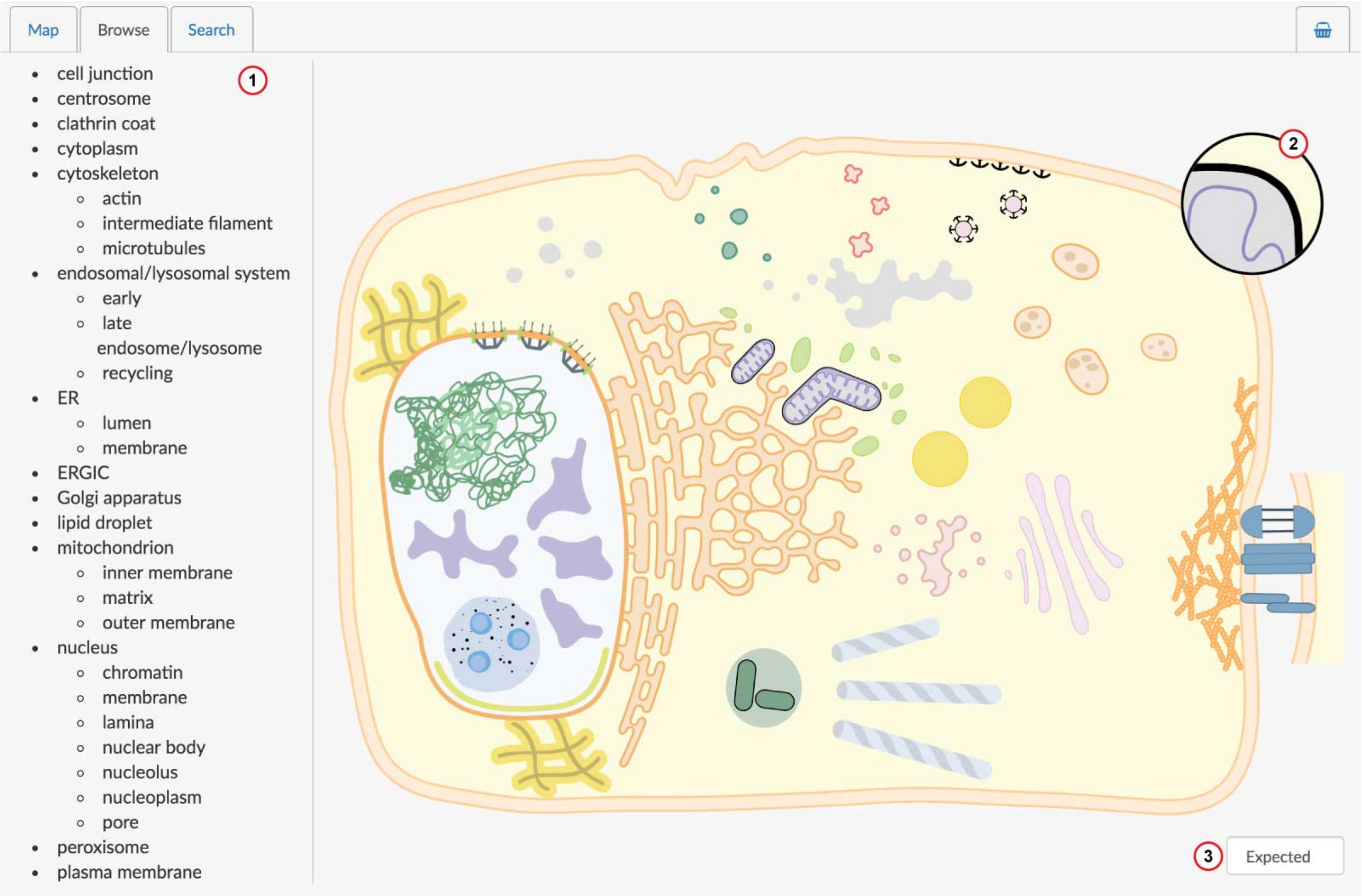


**Cartoon representation of the cell for viewing profiled organelles and selecting reports.**

1. List of compartments profiled or assigned through SAFE or NMF analysis
2. Inset for selecting mitochondrial subcompartments
3. Switch to NMF or SAFE views showing the organelles that were successfully mapped using those analyses

Hovering over any organelle on the cartoon will display its name or hovering over the name in the list on the left will highlight the organelle in blue. You can click on an organelle in the cartoon or its name on the list to open its report.

By default the cartoon will display the "Expected" view. From here users can navigate to the bait lists used to profile each organelle. From the dropdown menu at the bottom right you can switch to "NMF" or "SAFE" views showing the organelles that were successfully mapped using those approaches. If an organelle was not mapped for a particular approach, it will be greyed out in the cartoon.

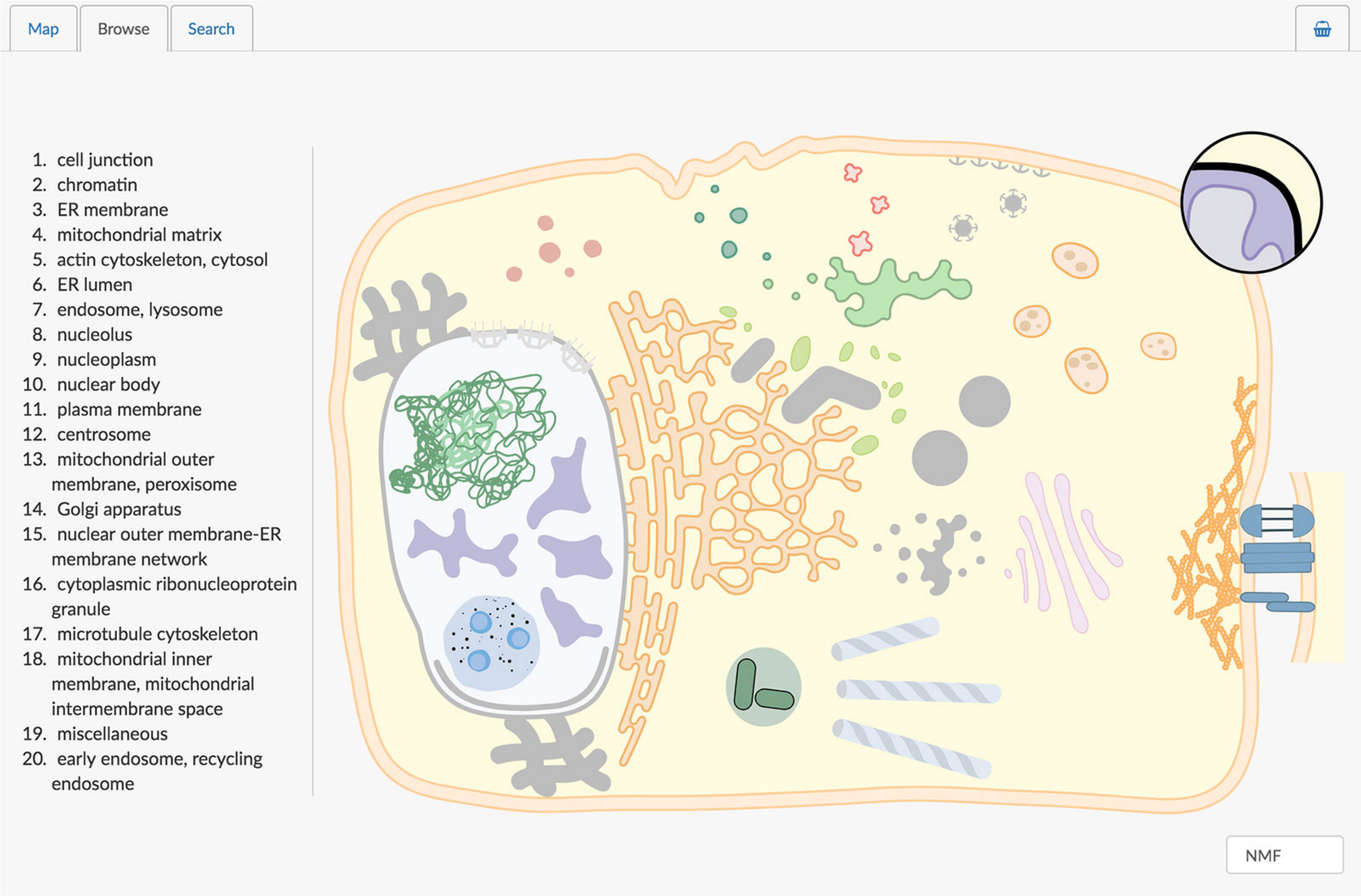


**The intermediate filament compartment was not identified using NMF and it is greyed out on the cartoon.**

#### Notes:

On the Expected view there are two organelles greyed out: cytoplasmic ribonucleoprotein complex and endosome. Although we did not explicitly select baits to profile these compartments, we were still able to map them using NMF and/or SAFE, hence they are shown on the cartoon. On both the NMF and SAFE views the mitochondria are greyed out. For these approaches we were able to assign preys to mitochondrial subcompartments and these can be selected using the inset in the top right.

### EXPLORE

#### SEARCH

The **Search** tab can be used to search for genes profiled as baits or detected as preys. By default, only preys passing an FDR cutoff of 0.01 will be displayed. Click the advanced options button + to change the significance cutoff (FDR), or to filter by cell type or publication (only one cell type and publication are currently available).

The "Baits" table will display any bait we profiled matching the search term, including the cell type it was profiled in and the publication associated with the data.

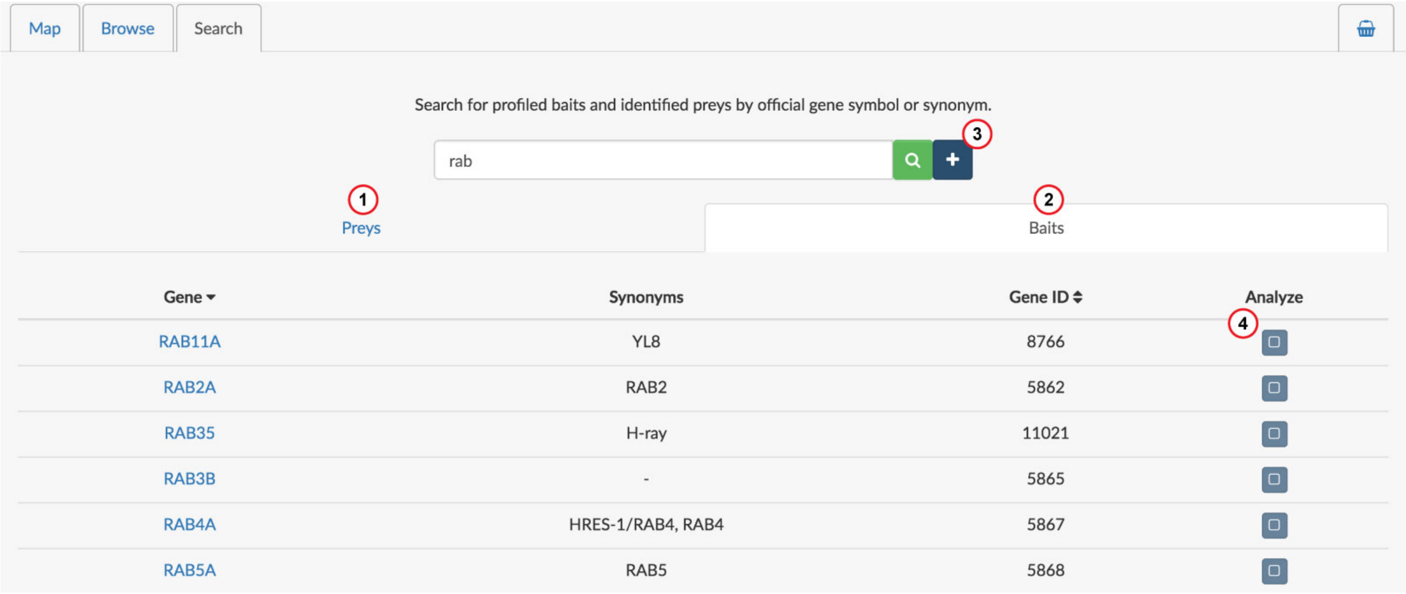


**Baits matching search term**

1. Preys matching search term
2. Baits matching search term
3. Advanced filtering options
4. Add bait to the analysis basket

The "Preys" table will show any preys matching the search term, along with the bait(s) they were seen with, the cell type, associated publication, spectral count (SC), average probability score (AvgP) and FDR. When a prey is identified with multiple baits, all of those instances will be reported along with the corresponding bait.

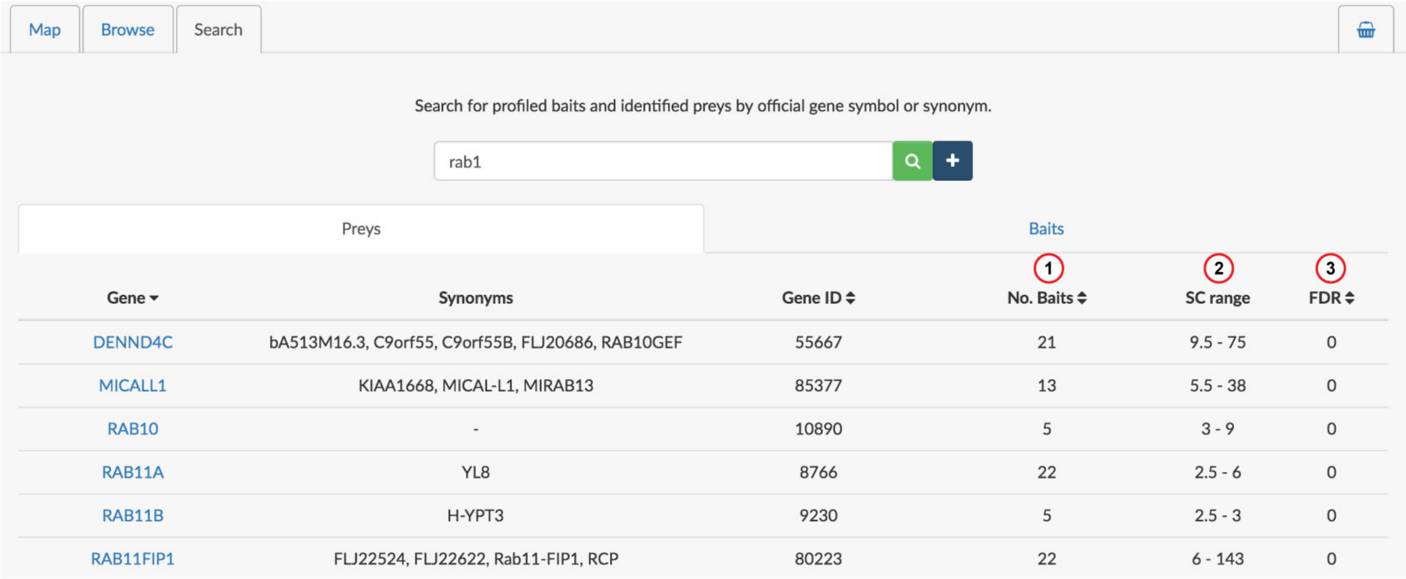


| Gene | Synonyms | Gene ID | No. Baits | SC range | FDR |
|---|---|---|---|---|---|
| DENND4C | bA513M16.3, C9orf55, C9orf55B, FLJ20686, RAB10GEF | 55667 | 21 | 9.5 - 75 | 0 |
| MICALL1 | KIAA1668, MICAL-L1, MIRAB13 | 85377 | 13 | 5.5 - 38 | 0 |
| RAB10 | - | 10890 | 5 | 3 - 9 | 0 |
| RAB11A | YL8 | 8766 | 22 | 2.5 - 6 | 0 |
| RAB11B | H-YPT3 | 9230 | 5 | 2.5 - 3 | 0 |
| RAB11FIP1 | FLJ22524, FLJ22622, Rab11-FIP1, RCP | 80223 | 22 | 6 - 143 | 0 |

**Preys matching search term**

1. The number of baits the prey was seen with at the specified FDR threshold (default 0.01)
2. Range of spectral counts for the prey. This is calculated across the baits it was detected with at the specified FDR threshold (default 0.01).
3. Best (lowest) false discovery rate (FDR)

Click on any gene name to open its report. You can also Control-click (Command ⌘ for Mac users) on a name to open the report without navigating to it (i.e. to open a report but remain on the search page).

### EXPLORE

#### REPORT

The **Reports** tab is used for viewing bait, prey and organelle reports. Tabs will be sorted by type and alphabetically within a type. The different report types will have tabs colour coded as indicated below:

Bait Organelle Prey

##### Organelle

Organelle reports can be of three types: "expected", "NMF" or "SAFE". An "expected" organelle report will show all baits profiled in the cell map with that expected localization. An "NMF" or "SAFE" organelle report will show all preys that localize to that compartment, along with the enriched GO terms, protein domains and motifs (if any).

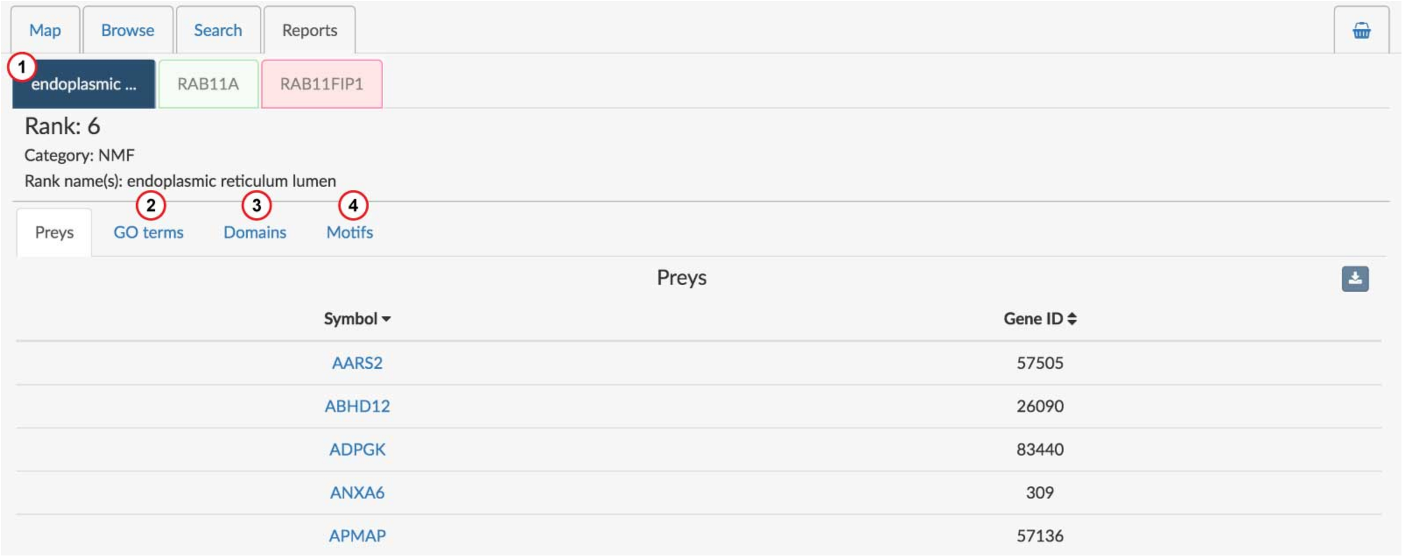


**NMF organelle report for the endoplasmic reticulum lumen**

1. Organelle report tab
2. Enriched GO terms
3. Enriched protein domains
4. Enriched protein motifs

##### Bait

Bait reports display basic information about the bait, its expected subcellular localization, links to external resources and preys that were detected with it. From the **Localization** tab you can find a list of enriched GO terms generated from preys detected with the bait and any immunofluorescence images that were used to validate bait localization. Data that is displayed by default on the prey table must pass certain filters that can be viewed and adjusted by clicking the **Filters** button.

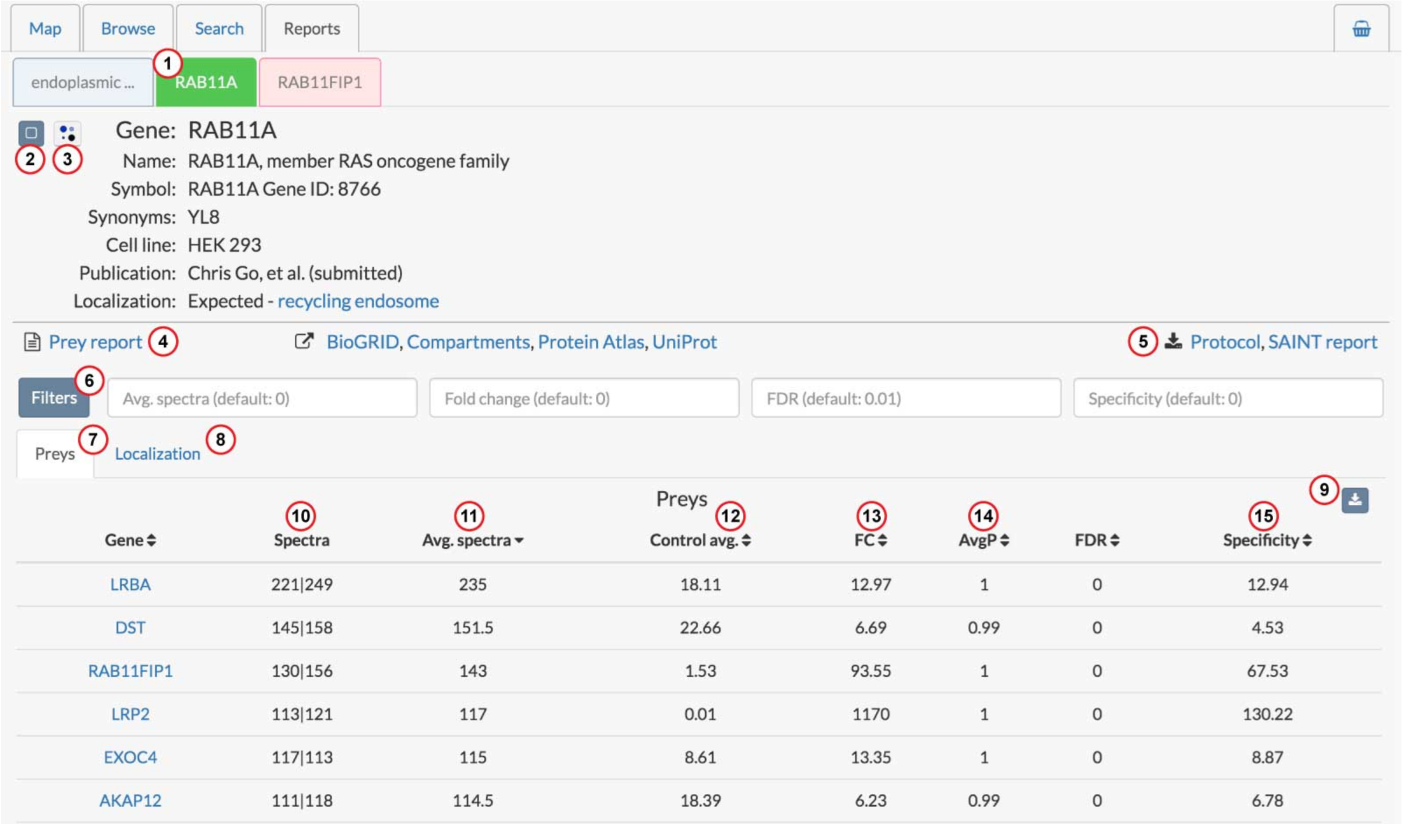


**Bait report for RAB11A**

1. Bait report tab
2. Add bait to analysis basket
3. Compare bait with any other open bait reports
4. View the prey report for the bait (only present when the bait was also seen as a prey)
5. Download the protocol used for profiling the bait or the full SAINT report
6. Filter the list of preys
7. List of preys identified by the bait
8. Localization information for the bait
9. Download the prey list
10. Spectral count across the two biological replicates
11. Average spectra of the two biological replicates
12. Average spectra across all controls
13. Fold-change of average spectra in bait sample relative to control samples
14. SAINT score
15. Specificity score, measured as the fold enrichment for the prey relative to the entire dataset

#### Prey

Prey reports display basic gene information, links to external resources, baits that detected the prey, other highly correlated preys, and both NMF and SAFE localization information. Data displayed on both the bait and prey tables can be adjusted using the "Filters" button.

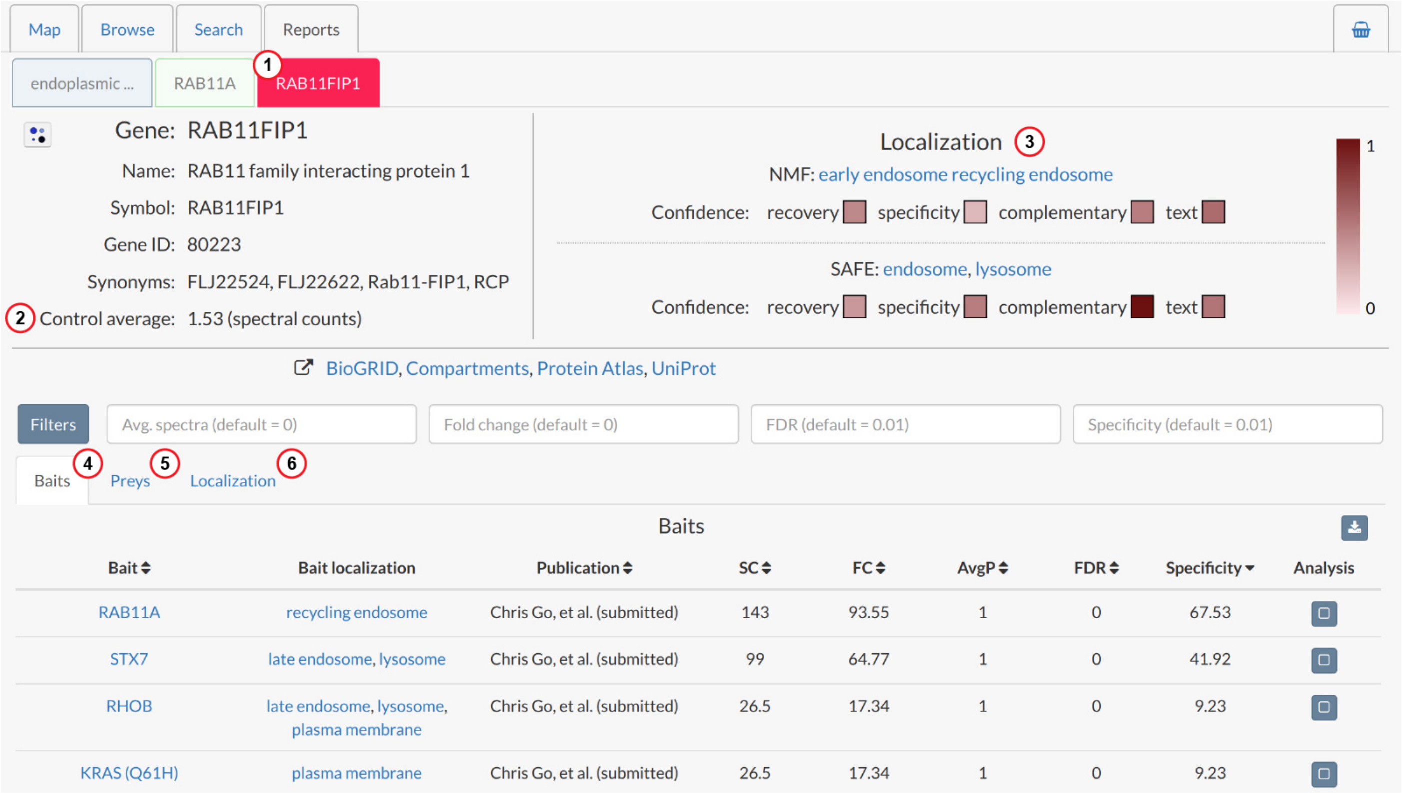


**Prey report for RAB11FIP1**

1. Prey report tab
2. Average spectra for the prey in control samples
3. Localization and confidence metrics
4. List of baits that identified this prey
5. List of correlated preys
6. Localization information

The localization panel indicates the predicted NMF and SAFE localizations along with metrics for gauging the confidence of these predictions. The confidence metrics are on a scale of 0-1 with 1 being the highest confidence level.

- recovery: the proportion of baits localizing to the predicted compartment that the prey was detected with at a 1% FDR. If a prey localizes to the ER lumen and was detected with 6/8 ER lumen baits, it would get a score of 0.75.
- specificity: the proportion of baits the prey was detected with at a 1% FDR that localize to the predicted compartment. A prey localized to the nucleus that was detected with 10 baits, seven of which are nuclear, would get a score of 0.7. If it was detected with 100 baits and only seven are nuclear, it would get a score of 0.07.
- complementary: agreement with other prediction techniques. Using microscopy-based predictions from the Human Protein Atlas and combined fractionation predictions from the studies of Itzhak, et. al. and

Christoforou, et. al. , this metric reports the agreement of our predictions with these two approaches. A score of 1 indicates agreement with both, a score of 0.5 indicates agreement with one and 0 with neither.

- text: literature evidence from the Compartments database text mining score. This database provides a confidence score for GO compartment terms for genes based on a gene's co-occurrence with subcellular localization terms in Medline abstracts. The score for our predicted compartment is extracted and converted to a 0-1 scale.

#### Comparisons

If you have multiple bait or prey reports open, you can compare them as a dot plot using the button located at the top left of the report page. Simply click the button, choose which baits or preys you want to compare and hit submit.

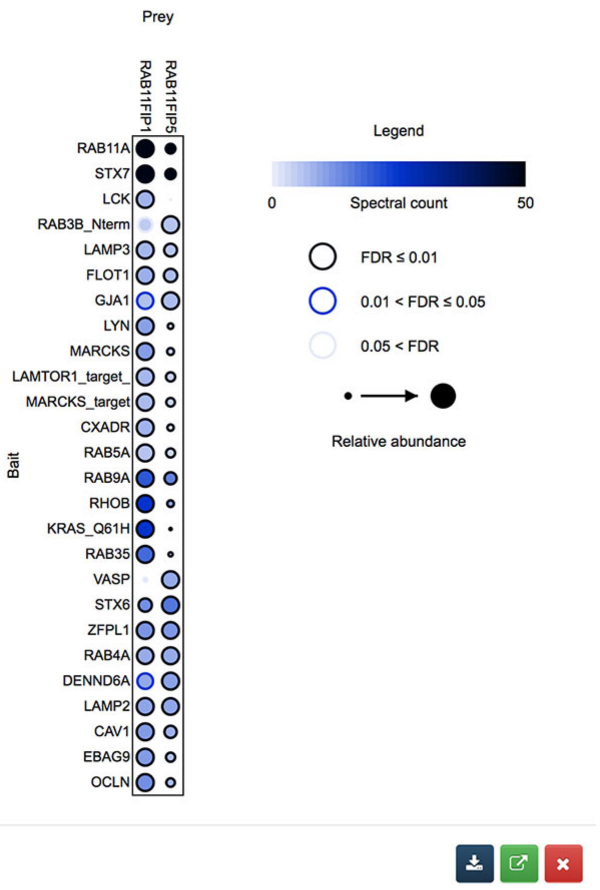


**Dot plot comparison of RAB11FIP1 and RAB11FIP5**

The image can be downloaded from by clicking the button or it can be opened at ProHits-viz by clicking the button.

#### Downloads

You can download any table on report tabs using the download button:

### EXPLORE

#### ANALYSIS-BASKET

Baits can be selected for analysis anywhere on search and report pages where you see a ▢ button. From the **Analysis basket** tab you can then compare the selected baits as a dot plot or view the baits and preys as a network. The resulting images can be downloaded as an SVG or opened at ProHits-viz. You can also download a SAINT report for the baits in the basket by clicking the SAINT button. The SAINT report can be used to perform your own analysis at ProHits-viz or for generating networks with Cytoscape.

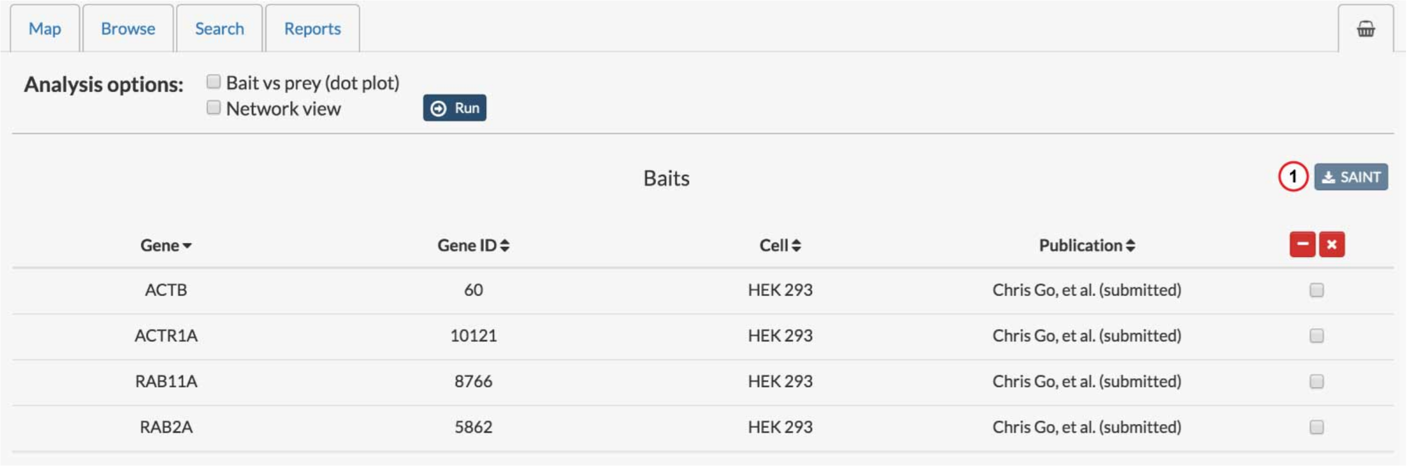


**Baits selected for comparison in the analysis basket**

(1) Download the SAINT report for all baits in the basket

For details on the types of images produced, please see:

- prohits-viz.lunenfeld.ca
- Dot plot description - PMID:25422071
- ProHits-viz - PMID:28661499
- Cytoscape - PMID:14597658

#### ANALYSIS

Users can compare their own BioID data against our entire database or against a set of baits localizing to a particular organelle. Comparing against the entire database can help to localize a bait, while more specific organelle-targeted comparisons can identify preys that are specific to the queried bait.

We provide this service to give others a context in which to interpret their own data. Users with small datasets may find this particularly helpful as it can prevent over-interpretation of results.

### ANALYSIS

#### INPUT FILE

Explicit support is provided for data files output from SAINT and CRAPome/REPRINT. Datasets from other tools or pipelines can be input by selecting the "Generic" option. Files must be in tabular format as tab-delimited text. At a minimum, the file must contain four columns specifying the bait, prey, abundance measure (spectral count or intensity) and a confidence metric (e.g. FDR). Sample input files are available for download and contain BioID data published in PMID:24255178.

For detailed information on tools that generate compatible input, see the references and links below.

##### Publications:

- SAINT - PMID:21131968
- SAINTexpress - PMID:24513533
- SAINT-MS1 - PMID:22352807
- ProHits - PMID:20944583
- ProHits Protocol - PMID:22948730
- ProHits 4.0 - PMID:27132685
- CRAPome - PMID:23921808

##### Sites:

- saint-apms.sourceforge.net
- ProHits
- REPRINT
- GalaxyP

### ANALYSIS

#### SETTINGS

After selecting the file for upload and specifying the file type, hit the process button and follow the instructions. You will need to specify the columns to use for analysis, whether or not control subtraction should be performed and the score details. For control subtraction, you can specify a column with abundance values found in control samples and the average of these will be subtracted from your prey value prior to analysis. These control values must be supplied as a pipe-separated list (see the samples files for examples).

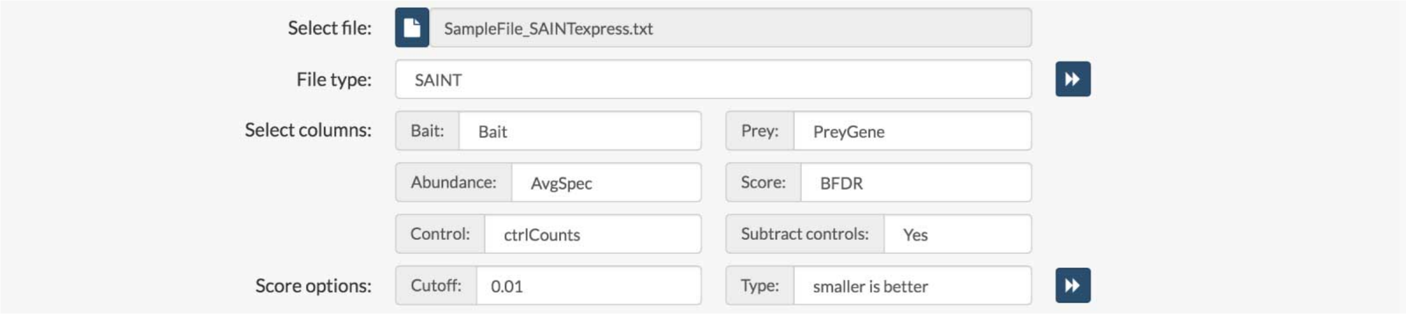


**Input file settings for analysis**

For the selected score, you must specify what cutoff should be used to identify significant interactors in your dataset and how the score works, i.e is a lower score better than a higher one, or vice versa. If you have output generated by SAINT or the CRAPome, options will be automatically selected for you although they can be adjusted as needed.

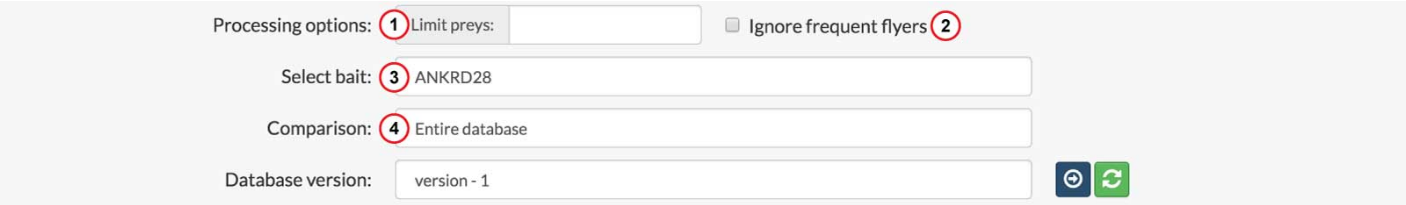


**Processing options for analysis**

1. The maximum number of preys to use in analysis
2. Remove "frequent flyers" from the list of preys
3. Select a specific bait from the uploaded data to be compared
4. Perform comparison against the entire Human Cell Map database, or select individual organelles/compartments

By default all preys passing the specified cutoff will be used for comparisons against the cell map. However, the number of preys can be limited and frequent flyers can also be ignored. If you choose to limit the number of preys to use, preys will be sorted by abundance and only the top preys up to and including your limit will be used for comparisons (both for your bait and the cell map baits). Frequent flyers are preys that are commonly seen with baits in the cell map and hence are less informative. If you choose to ignore frequent flyers, these preys will be ignored when performing comparisons. The list of frequent flyers is available for download. If both options are selected, frequent flyers will be removed first before the top preys are selected.

You can also choose whether a comparison should be done against the entire cell map or a specific organelle. If you choose to compare against one or more organelles, only baits that localize to those organelles will be used for comparisons.

### ANALYSIS

#### REPORT

The heat map on the report displays the prey spectral count/intensity for the query bait and up to its ten most similar baits from the cell map, as determined using the Jaccard distance. The cell map baits are sorted left to right from most to least similar. There is no cutoff for this comparison, meaning that even if your bait is not very similar to anything in the cell map, the most similar baits will still be shown. Always inspect the heat map to assess the degree of similarity.

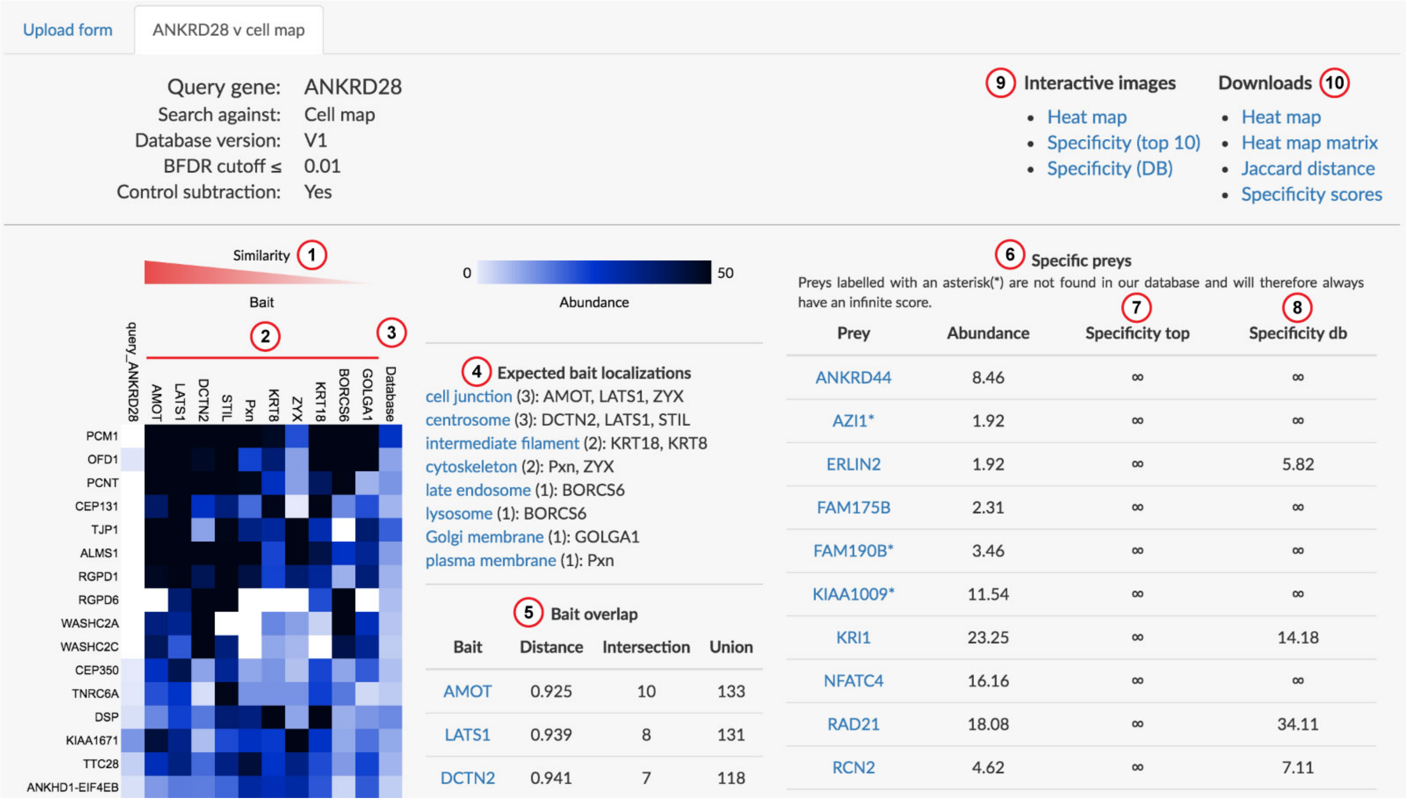


**Analysis report for a query bait (ANKRD28)**

1. Baits from the cell map are sorted from most similar to least similar as calculated by the Jaccard distance
2. Ten most similar baits to the query in the cell map
3. The average spectral count for each prey averaged across all baits in the cell map
4. Expected localizations of the ten most similar baits
5. Overlap/similarity metrics between the query bait and the top ten most similar baits in the cell map. The distance is the Jaccard distance, with a score of 0 for complete prey overlap and 1 for no overlap. The intersection refers to the number of shared preys and the union refers to the combined number of preys between the query and the indicated bait.
6. The most specific preys for the query. The specificity score is calculated as the fold enrichment of a prey in the query relative to the average across the cell map baits used for the comparison
7. The specificity score calculated against the top ten most similar baits to the query
8. The specificity score calculated against all baits in the cell map

9. Open the heat map or specificity plots at ProHits-viz
10. Download tabular data for the analysis

The right-most column on the heat map displays the average spectral count for each prey across all baits in the database. This is included to give users an idea of the average baseline value detected across our samples. The right panels show information about baits most similar to the query bait, as well as the preys most specific to the queried bait.

### DOWNLOAD

When using information or data from the human cell map, please cite: Go et al. (submitted) Journal, Volume, Page numbers.

Data from the human cell map is licensed under a Creative Commons Attribution 4.0 International License license. If you are unsure whether or not your intended use is allowable under this license, please contact us at.

You can download the latest and previous versions of the database programmatically from:

```
https://cell-map.org/resources/downloads
```

See the downloads page for more.

### CITATION

When using information or data from the Cell Map, please cite: Go et al. (in preparation), "A proximity biotinylation map of a human cell".
