## Extended data figures for "A proximity-dependent biotinylation map of a human cell: an interactive web resource"

### Extended Data Figure 1

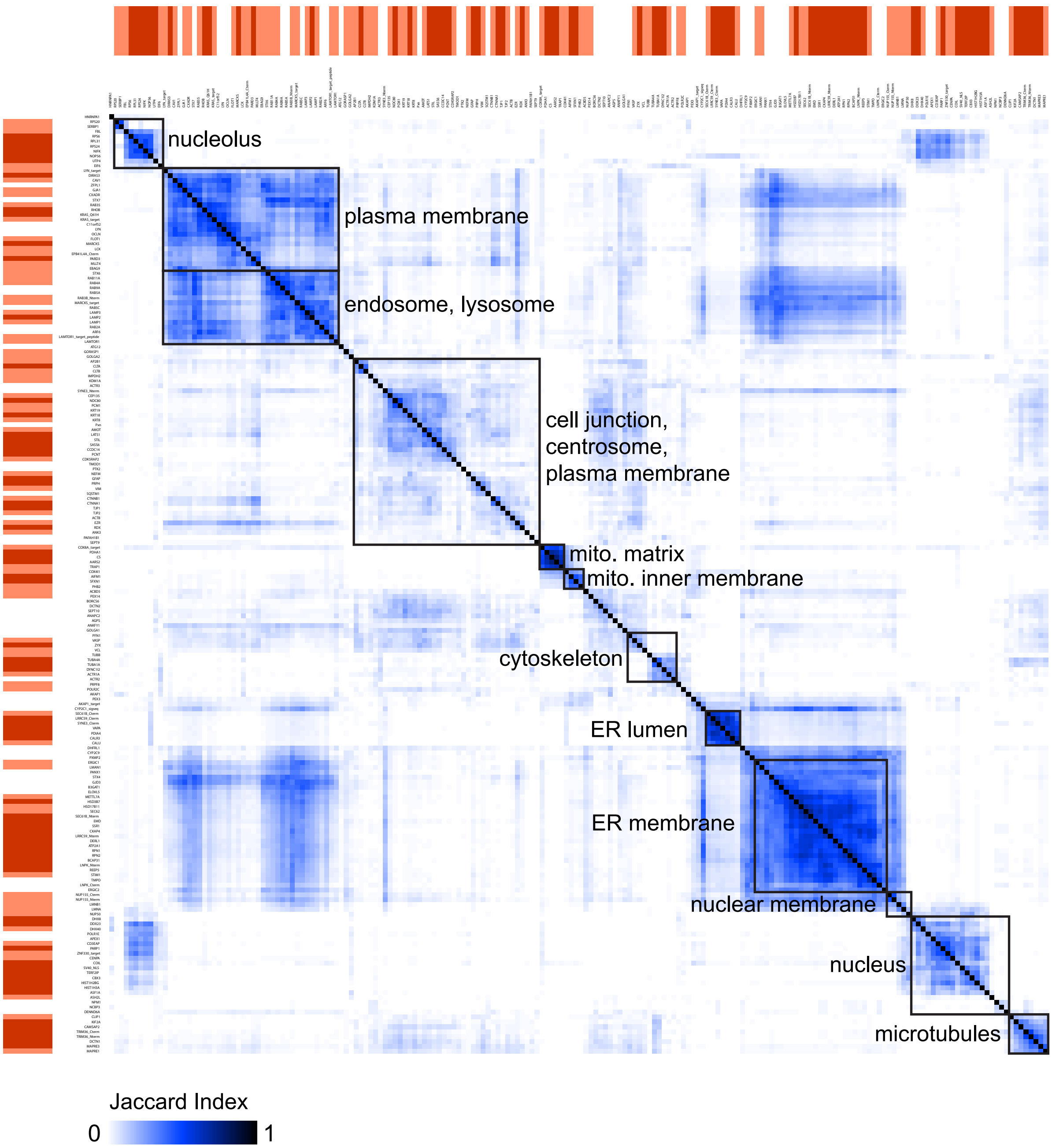

### Extended Data Figure 2

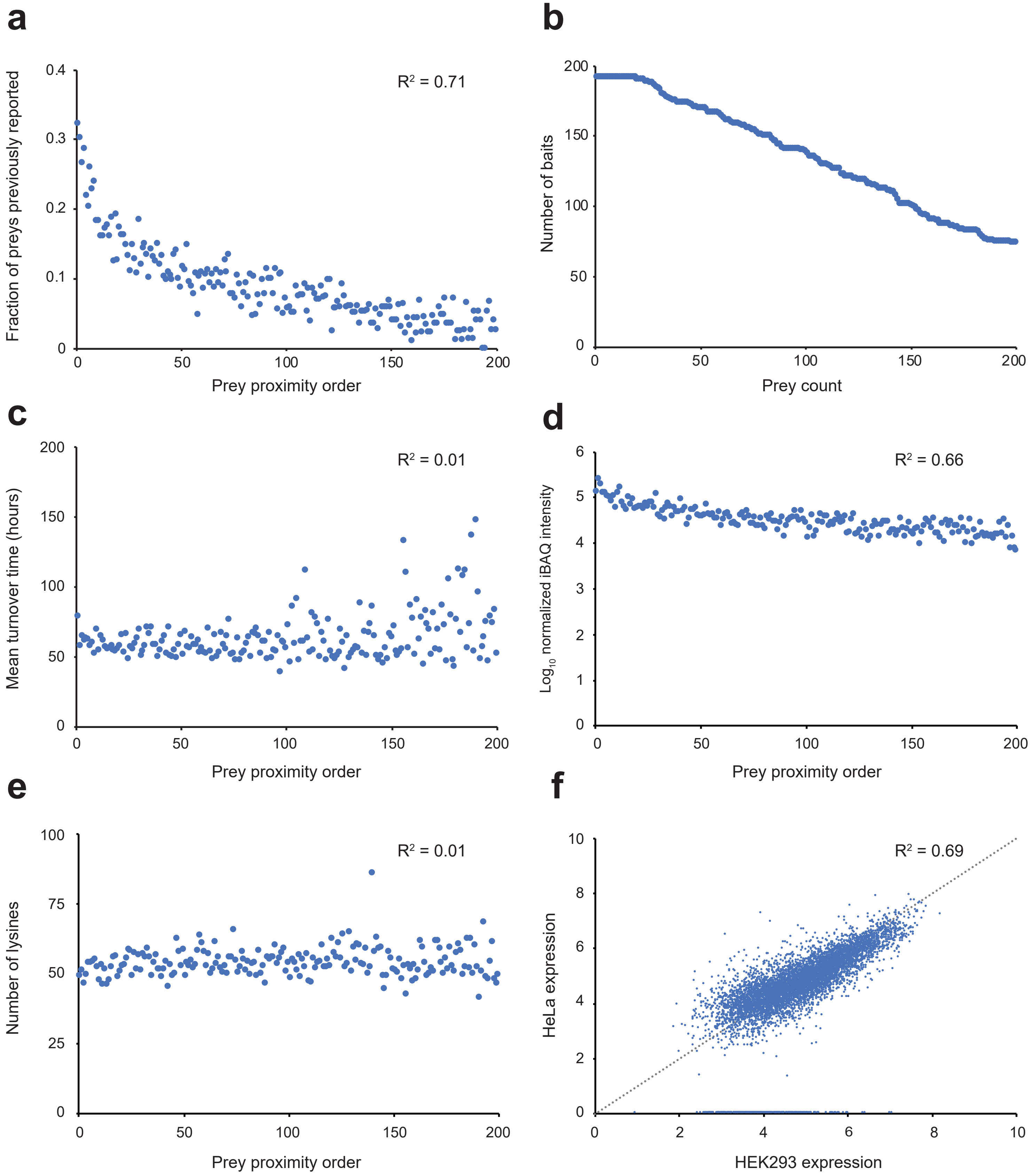

### Extended Data Figure 3

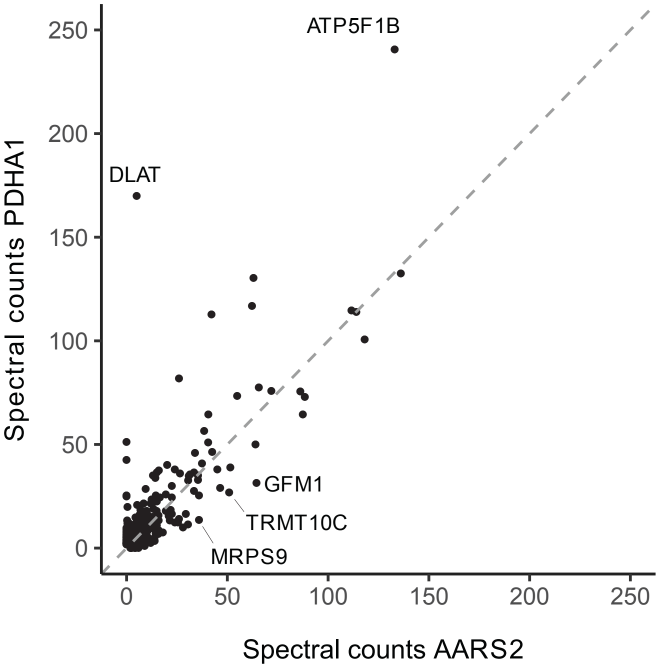

### Extended Data Figure 4

**a**

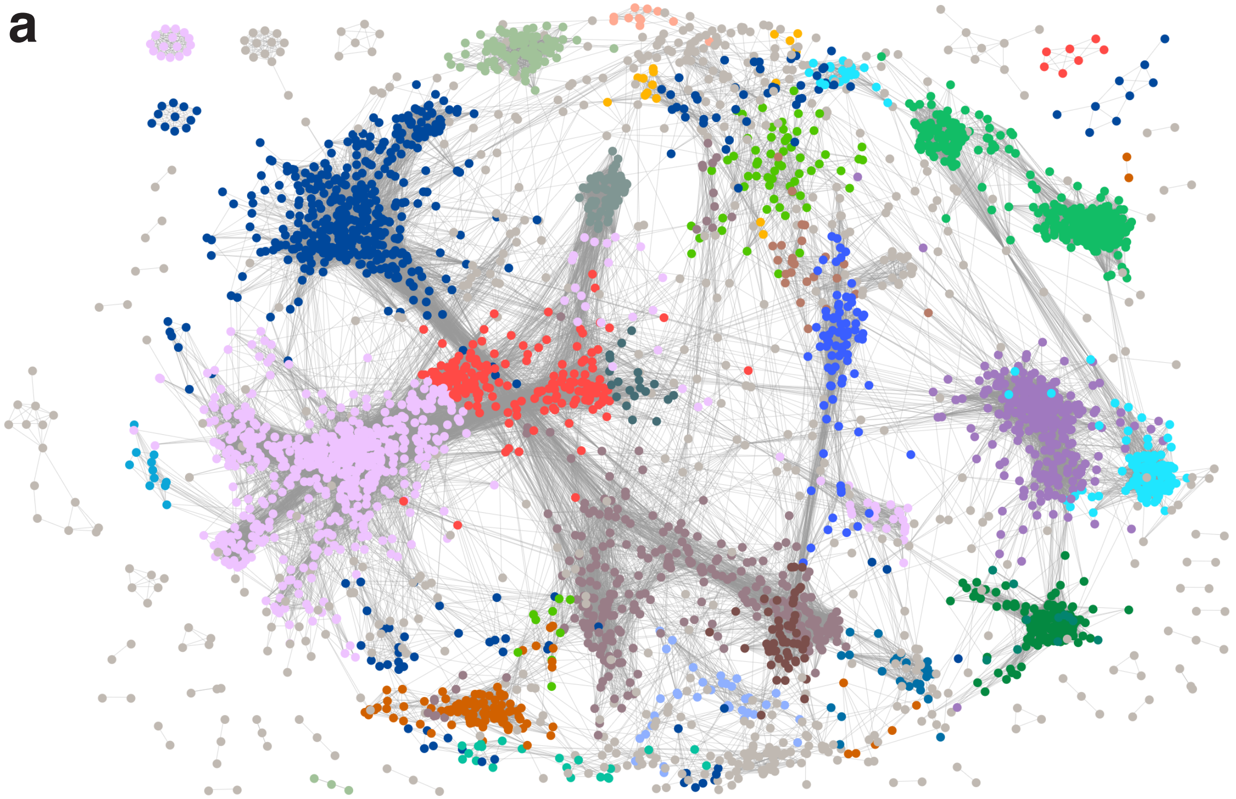

1. unknown
2. mitochondrial outer membrane, peroxisome
3. splicing speckles, nucleolus
4. ER lumen
5. spliceosomal complex I
6. clathrin coat
7. actin cytoskeleton
8. paraspeckles
9. Golgi apparatus
10. centrosome
11. chaperonin-containing T-complex, prefoldin complex
12. anaphase-promoting complex, kinesin complex
13. cytoplasmic ribonucleoprotein granule
14. centrosome enriched, P-body enriched
15. microtubule cytoskeleton
16. STRIPAK complex
17. Arp2/3 protein complex
18. endosome, lysosome
19. nuclear outer membrane-ER membrane network
20. cell junction, plasma membrane
21. mitochondrial inner membrane, mitochondrial intermembrane space, mitochondrial matrix
22. chromatin, nucleoplasm
23. spliceosomal complex II
24. nuclear pore

**b**

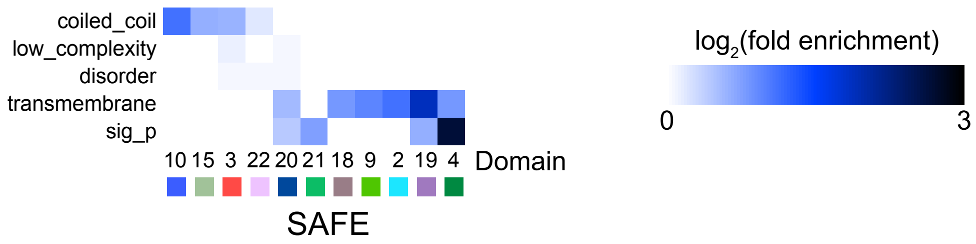

### Extended Data Figure 5

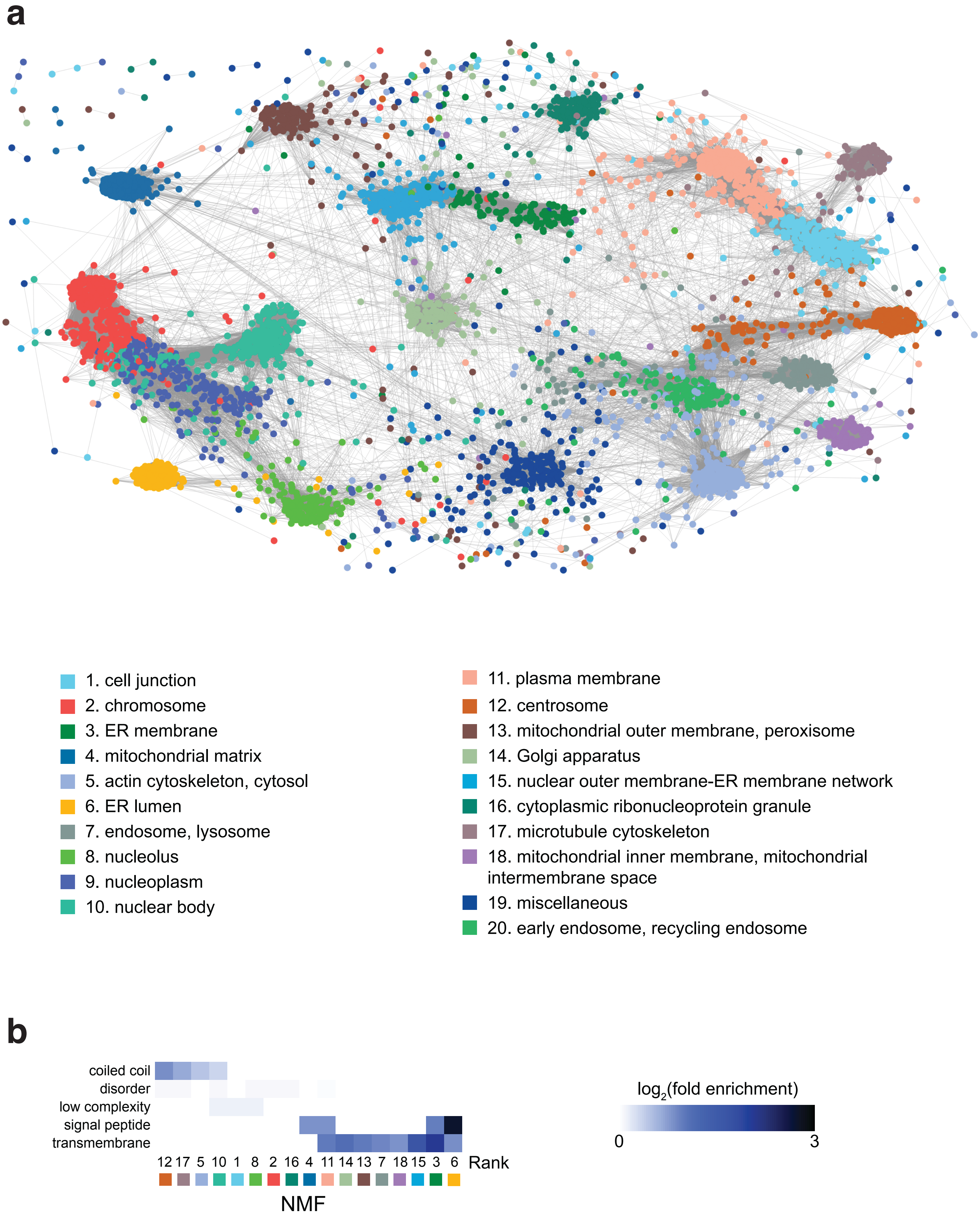

### Extended Data Figure 6

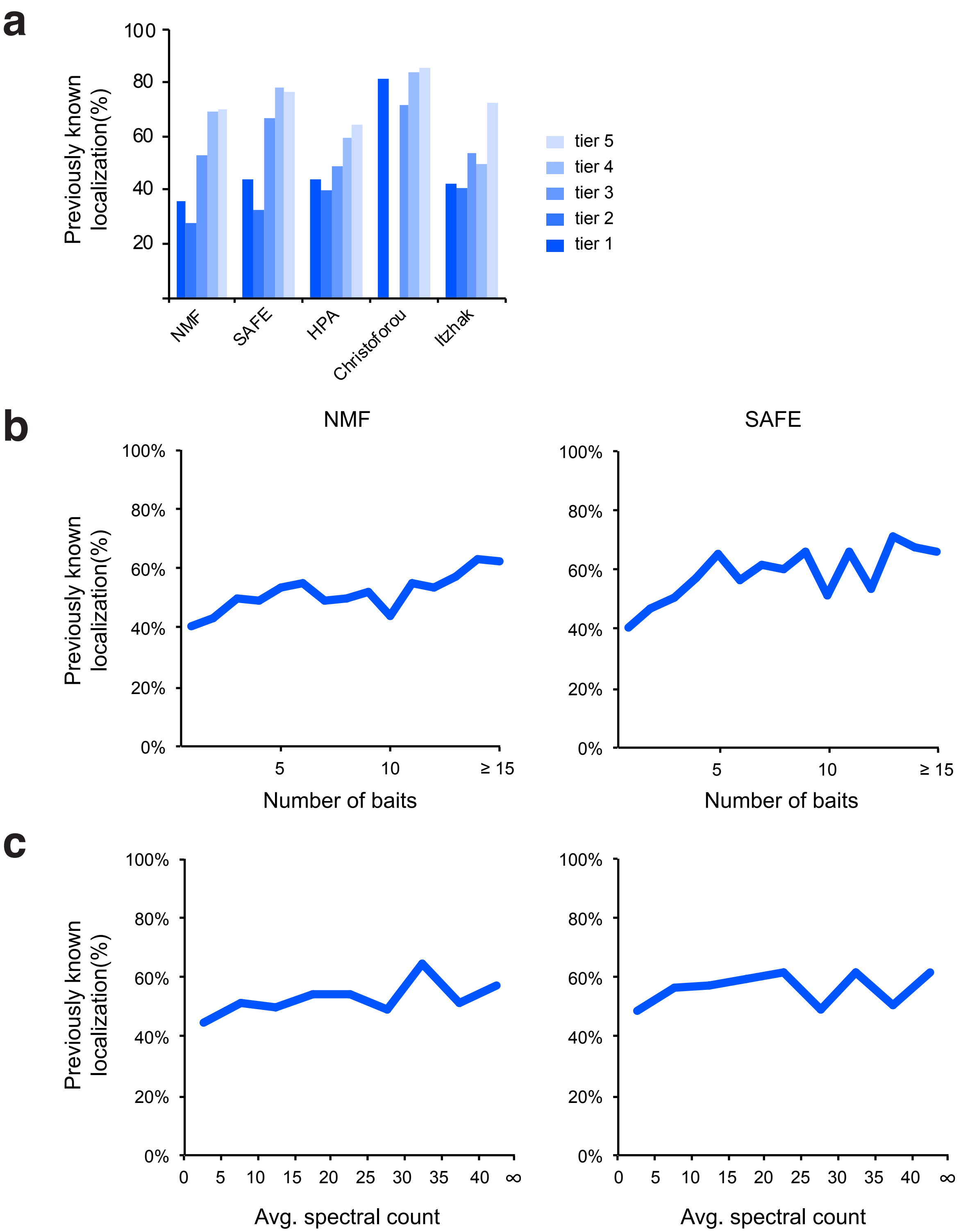

### Extended Data Figure 7

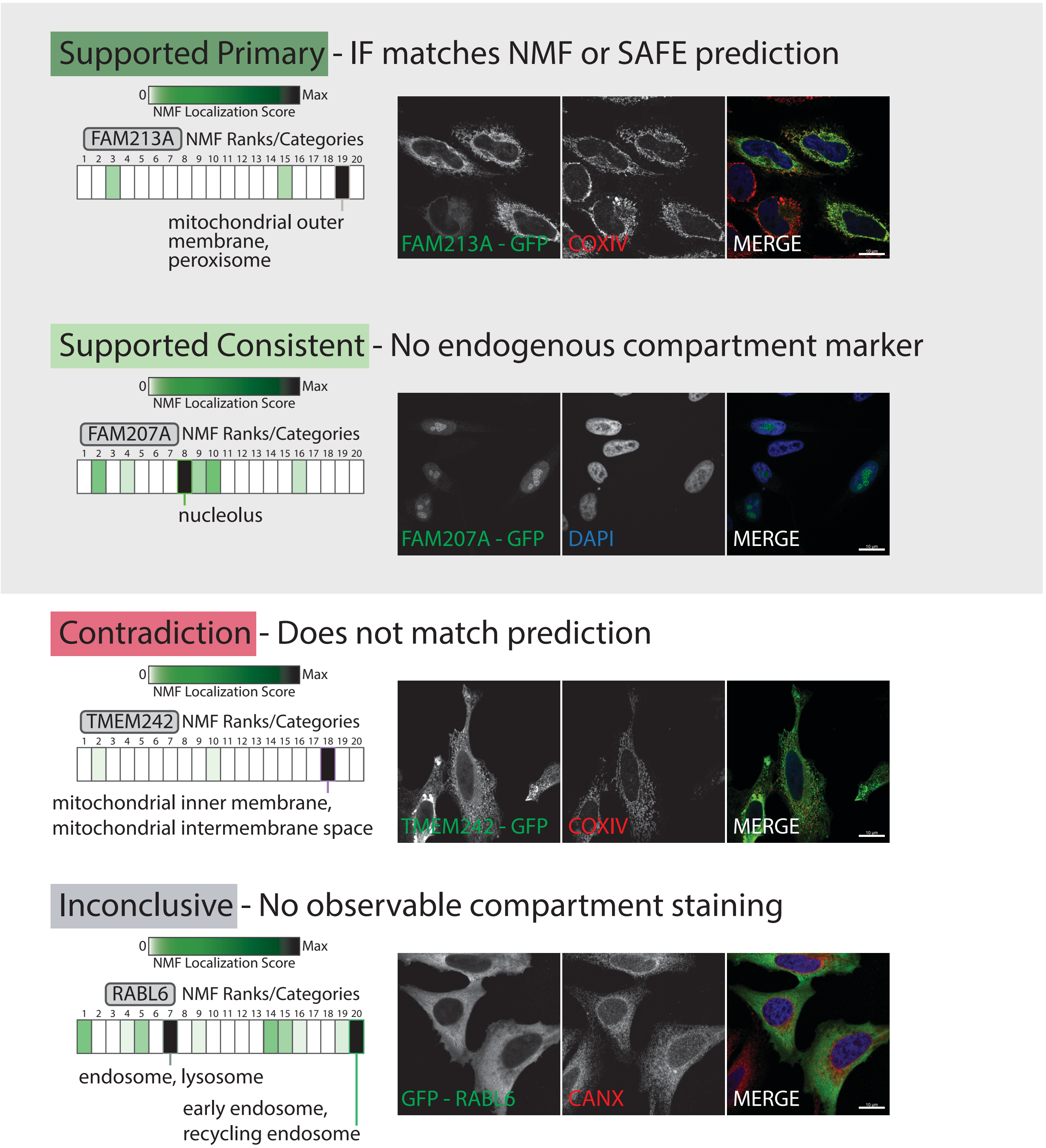

### Extended Data Figure 8

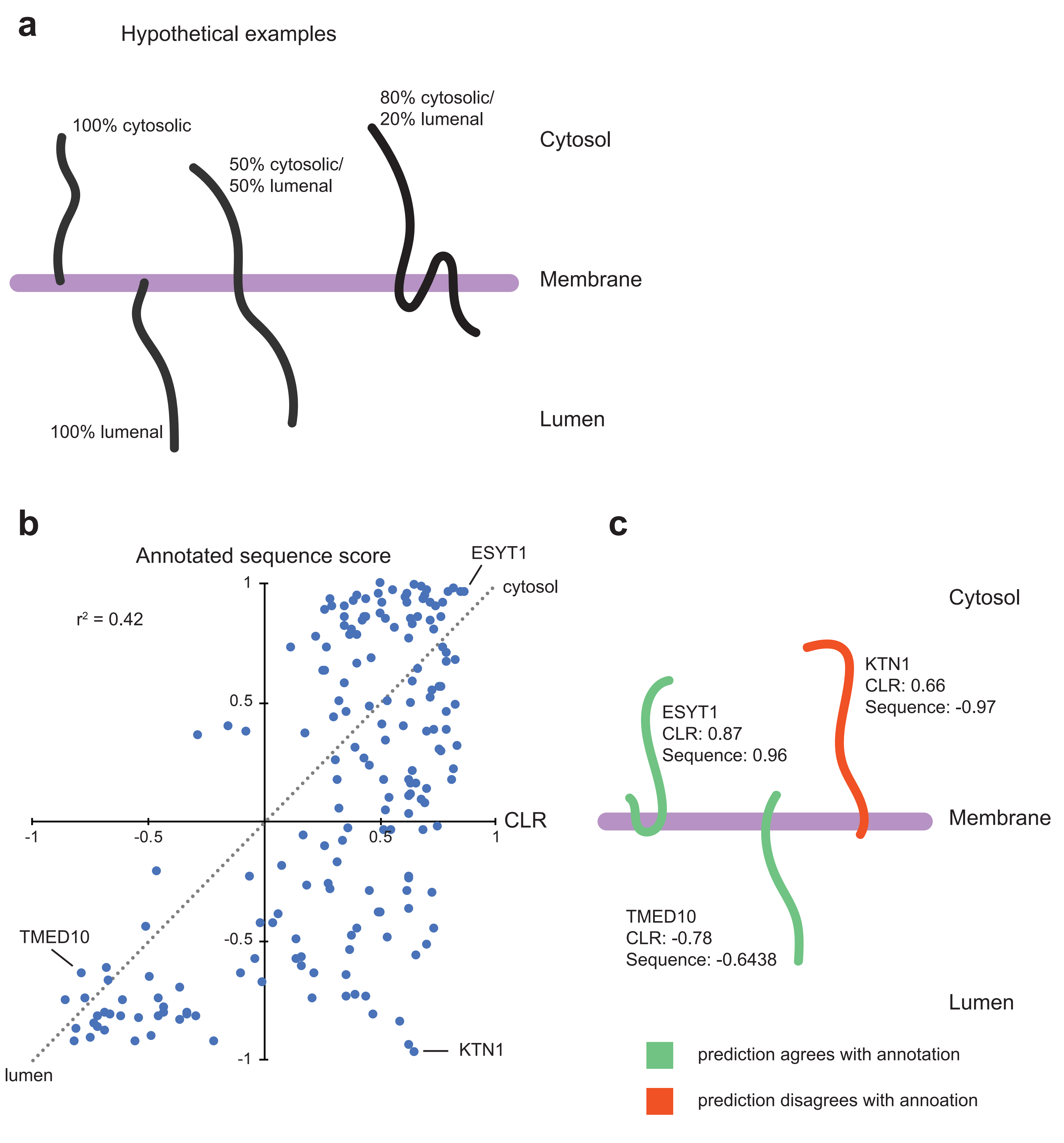

### Extended Data Figure 9

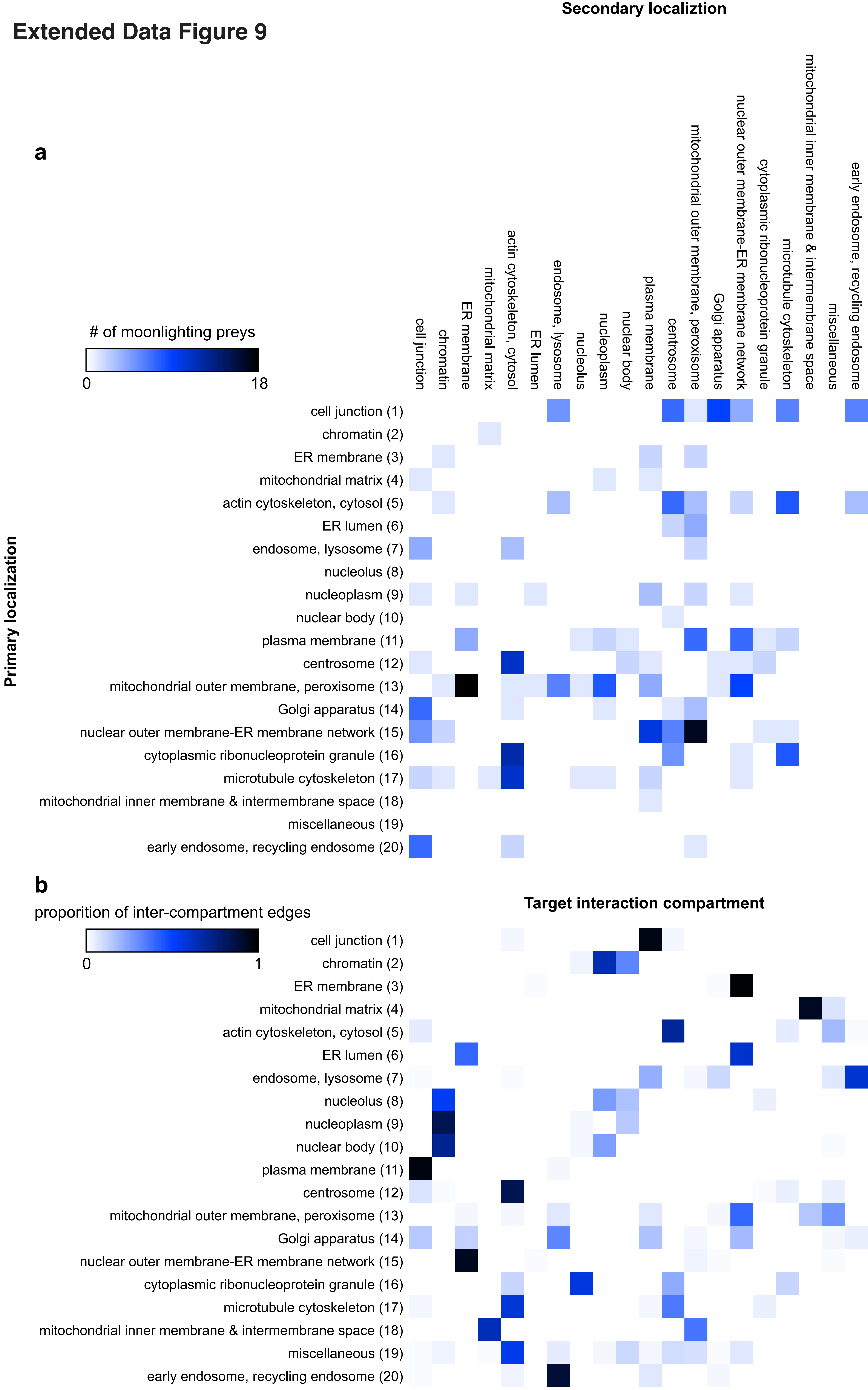

### Extended Data Figure 10

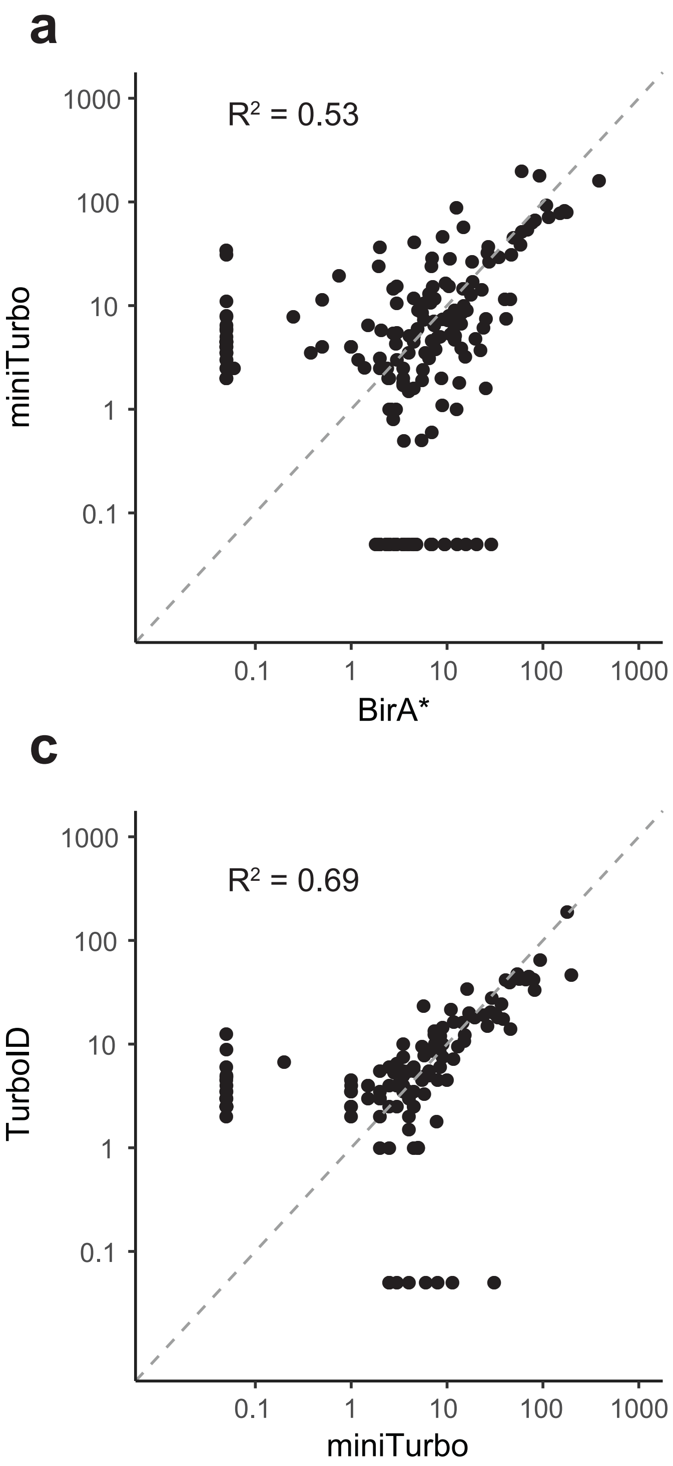

### Extended Data Figure 11

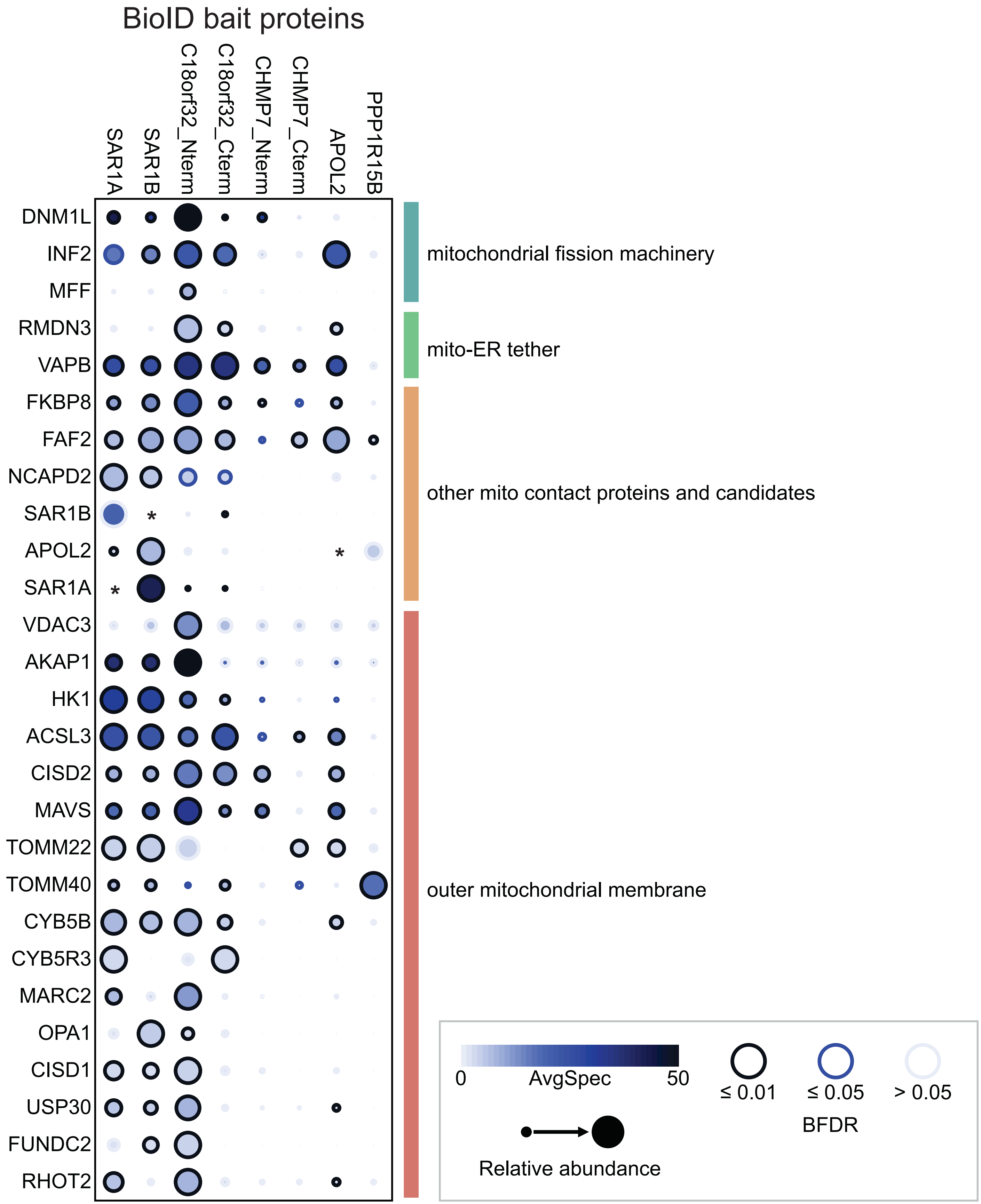

### Extended Data Figure 12

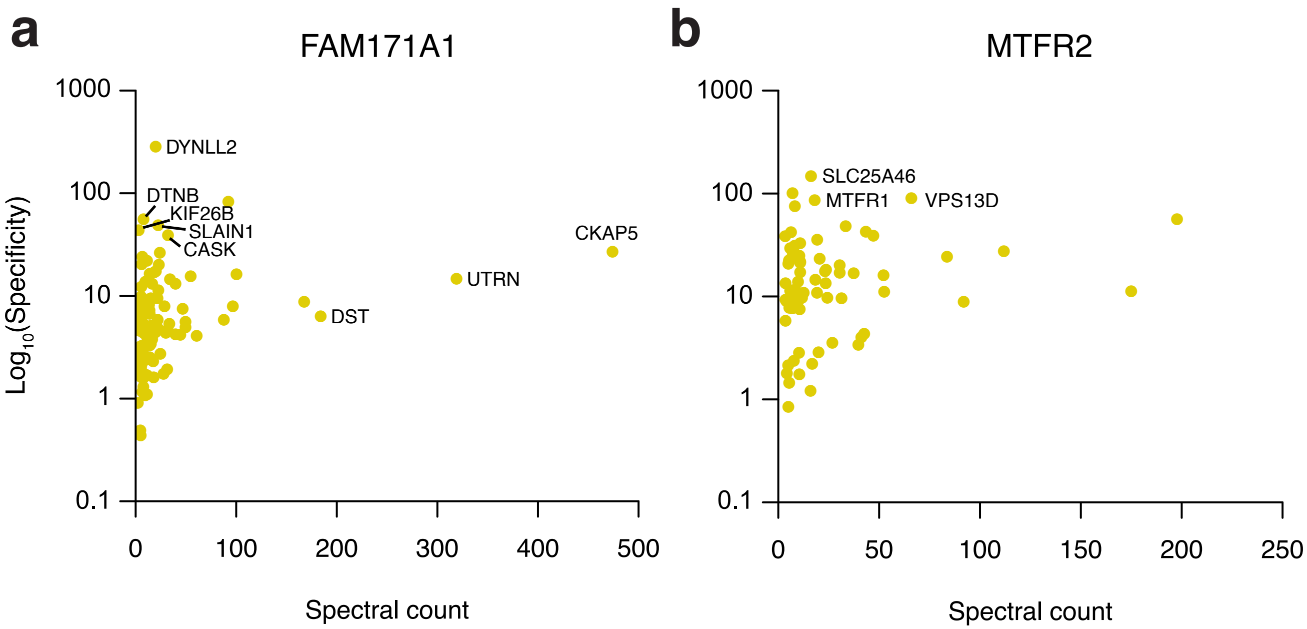

### Extended Data Figure 13

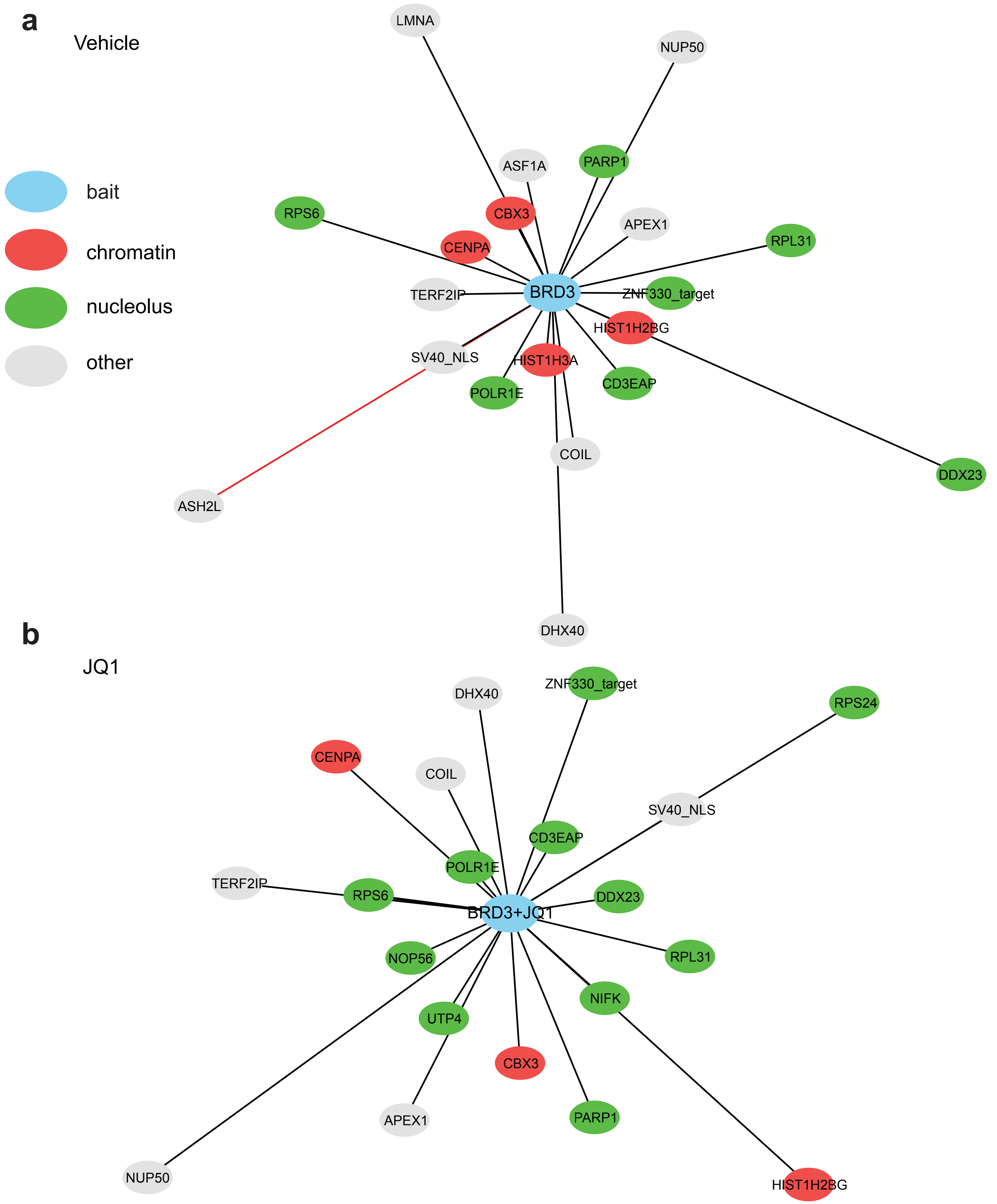

### Extended Data Figure 14

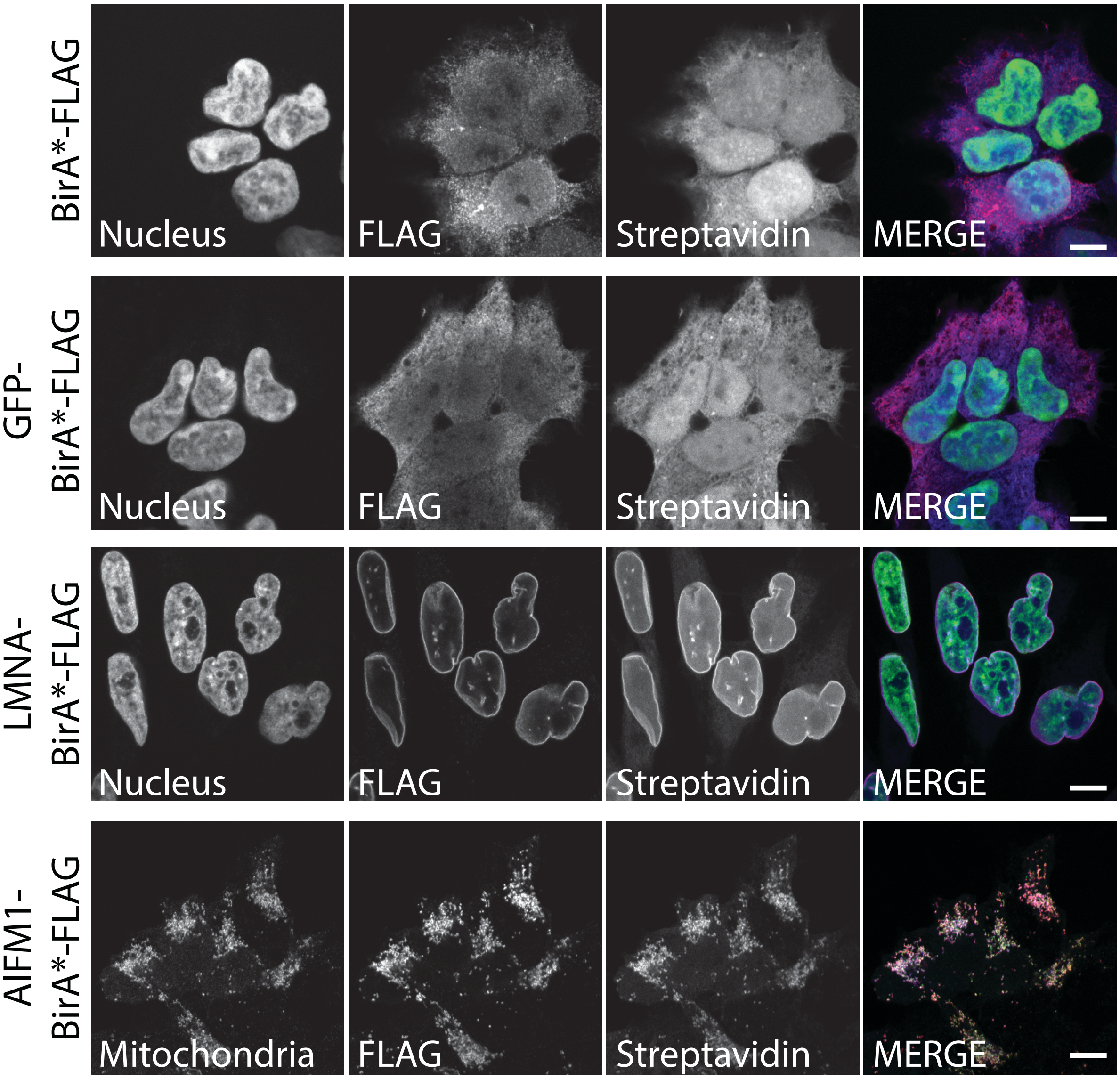
